# Sex-dependent dominance maintains migration supergene in rainbow trout

**DOI:** 10.1101/504621

**Authors:** Devon E. Pearse, Nicola J. Barson, Torfinn Nome, Guangtu Gao, Matthew A. Campbell, Alicia Abadía-Cardoso, Eric C. Anderson, David E. Rundio, Thomas H. Williams, Kerry A. Naish, Thomas Moen, Sixin Liu, Matthew Kent, David R. Minkley, Eric B. Rondeau, Marine S. O. Brieuc, Simen Rød Sandve, Michael R. Miller, Lucydalila Cedillo, Kobi Baruch, Alvaro G. Hernandez, Gil Ben-Zvi, Doron Shem-Tov, Omer Barad, Kirill Kuzishchin, John Carlos Garza, Steven T. Lindley, Ben F. Koop, Gary H. Thorgaard, Yniv Palti, Sigbjørn Lien

## Abstract

Traits with different fitness optima in males and females cause sexual conflict when they have a shared genetic basis. Heteromorphic sex chromosomes can resolve this conflict and protect sexually antagonistic polymorphisms but accumulate deleterious mutations. However, many taxa lack differentiated sex chromosomes, and how sexual conflict is resolved in these species is largely unknown. Here we present a chromosome-anchored genome assembly for rainbow trout (*Oncorhynchus mykiss*) and characterize a 56 Mb double-inversion supergene that mediates sex-specific migration through sex-dependent dominance, a mechanism that reduces sexual conflict. The double-inversion contains key photosensory, circadian rhythm, adiposity, and sexual differentiation genes and displays frequency clines associated with latitude and temperature, revealing environmental dependence. Our results constitute the first example of sex-dependent dominance across a large autosomal supergene, a novel mechanism for sexual conflict resolution capable of protecting polygenic sexually antagonistic variation while avoiding the homozygous lethality and deleterious mutation load of heteromorphic sex chromosomes.

---

Differential selection on male and female individuals results in sexual antagonism, with profound implications for genome evolution, adaptation, and the maintenance of fitness variation^1-4^. In the classical model of sex chromosome evolution, sexually antagonistic polymorphisms accumulate in linkage disequilibrium with the sex determining locus, driving selection for reduced recombination and differentiation of the sex chromosomes (X/Y, or Z/W), and the eventual degradation of the hemizygous chromosome through accumulation of deleterious mutations^5,6^. While this model explains sexual conflict resolution in some species, many taxa lack differentiated sex chromosomes^5^, and how sexually antagonistic variation is maintained in these species is largely unknown^4,7^. Additionally, recent theoretical work has predicted that sexual conflict may be better resolved by autosomal variation^8^, a prediction supported by genome-wide mapping of sexually antagonistic polymorphisms^3^. Thus, mechanisms that maintain sexual conflict polymorphisms on the autosomes must be common, yet the only known genetic mechanism for maintaining sexually antagonistic polymorphisms on autosomes, sex-dependent dominance^7,9^, has been observed only once for a single gene^4^. As a result, the mechanisms maintaining sexually antagonistic variation in the absence of differentiated sex chromosomes and their consequence for adaptive and genome evolution remain unresolved.

*Oncorhynchus mykiss* is a salmonid fish species that expresses two contrasting life-history strategies: Resident *rainbow trout* live entirely in freshwater, while anadromous *steelhead* migrate to the ocean to mature, returning to freshwater to reproduce. The decision to either mature early or delay maturation and migrate is a complex heritable trait, requiring the integration of internal (energy status/adiposity) and external (photoperiod and temperature) signals^10^. Survival of migratory juveniles to adulthood is very low, but fecundity of anadromous females can exceed that of resident females by an order of magnitude^10^. In contrast, males can mature early as freshwater residents, avoiding the high mortality associated with marine migration and employing a sneaker mating strategy to access paternity^10^. This sex-specific trade-off between reproduction and survival results in a greater frequency of anadromy in females^11^, and is predicted to drive sexual conflict over alternative migratory tactics with a shared genetic basis.

Autosomal inversion supergenes controlling alternative reproductive tactics and resembling sex chromosomes have been identified in some taxa, but suffer from homozygous lethality^12,13^ and concomitant chromosomal degradation^14,15^. Balancing selection can facilitate the evolution of novel dominance patterns and these, along with epistasis, are an important features of inversion polymorphism^16,17^. Typical of taxa with homomorphic sex chromosomes^5,8,18^, salmonids have undergone frequent sex chromosome turnover^19^, and sex-reversed males (XY females) have been suggested to occur^20,21^, both of which are predicted to limit divergence of the sex chromosomes^5,22^ and their ability to accumulate and protect sexual conflict polymorphisms^8,18^. Here we generate a chromosome-level genome assembly for rainbow trout and use it to identify a large autosomal inversion supergene influencing sexually antagonistic migratory tendency with sex-dependent dominance. Lacking homozygote lethality, this inversion complex provides a mechanism for maintaining polygenic sexually antagonistic variation whilst avoiding the deleterious mutation load accumulated by differentiated sex chromosomes.

## Rainbow trout genome reveals structural rearrangements

To characterise the genetic and genomic architecture of complex traits it is important to construct a high quality, chromosome-anchored whole genome sequence, a challenge in rainbow trout owing to the salmonid-specific whole genome duplication event (Ss4R) ~80-125 MYA^23-25^, and subsequent expansion of repetitive elements^24^. We created an assembly (<u>GCA_002163495.1</u>) containing 139,800 scaffolds with an N50 of 1.67 Mb and a total of 2.18 gigabases (Gb), representing more than 90% of the predicted length (2.4 Gb) of the rainbow trout genome^26^. High-density linkage mapping was used to order and orient scaffolds within linkage groups, producing 29 chromosome-length sequences containing 1.92 Gb (88.5%) of the genome assembly (Supplementary Information section 1). Most scaffolds not anchored to chromosomes consisted of repetitive sequences, which account for 57.1% of the rainbow trout genome (Supplementary Information section 4; Extended Data Figure S1), similar to the 59.9% previously reported for Atlantic salmon^24^. Annotation by the NCBI RefSeq pipeline predicted 53,383 genes, of which 42,884 are protein-coding (NCBI <u>*Oncorhynchus mykiss*</u> Annotation Release 100). Analysis of homeologous regions resulting from the salmon-specific duplication revealed 88 collinear blocks along 29 chromosomes (Figure 1; Supplementary Information section 2).

**Figure 1.**
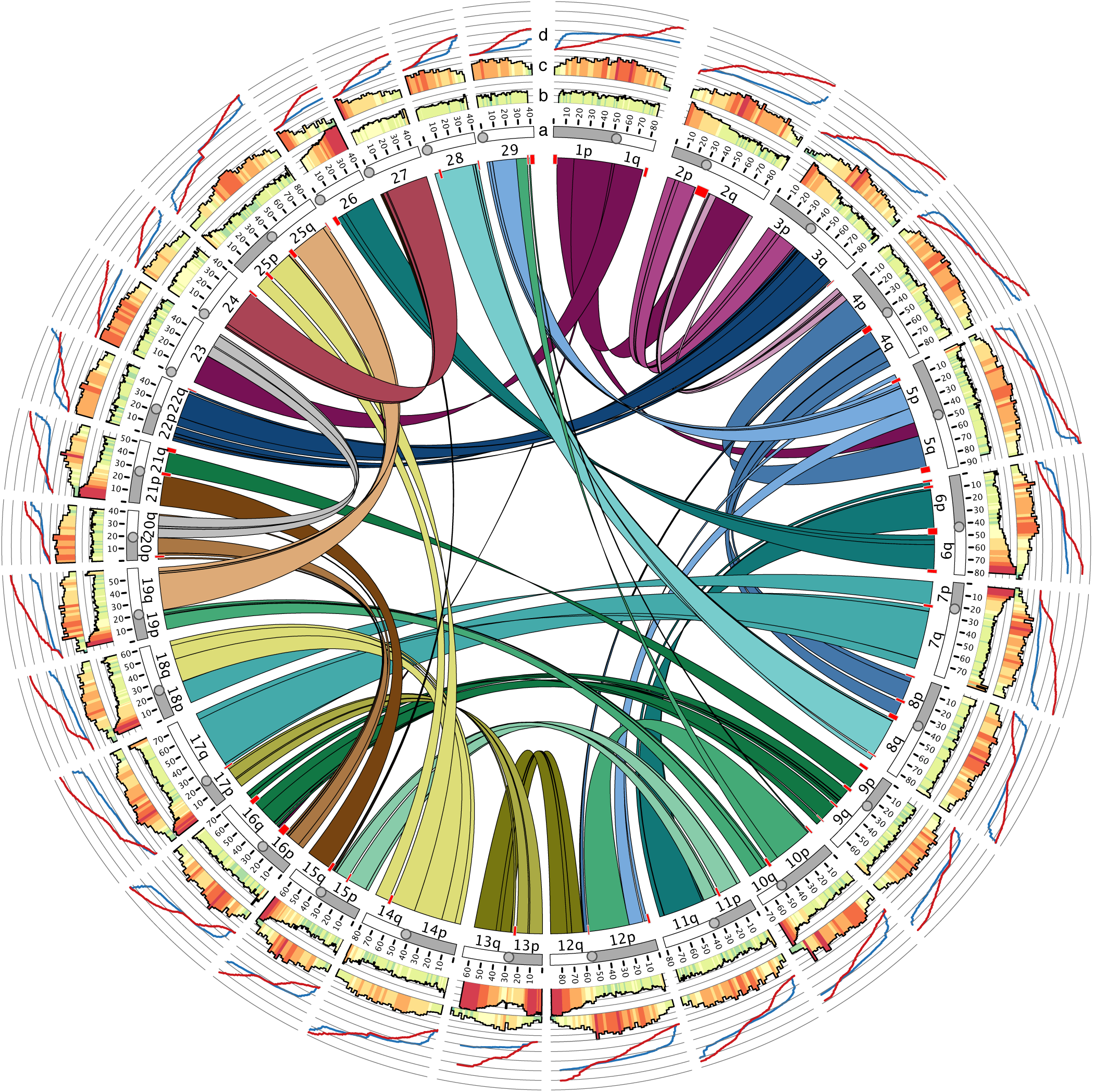
The duplicated rainbow trout genome, a-d inner to outer circles: A) Homeologous regions in the rainbow trout genome subdivided into 88 collinear blocks along 29 chromosomes. Red rectangles represent blocks of sequence without identifiable duplicated regions elsewhere in the genome. B) Genomic similarity (in 1DMb intervals) between duplicated regions. C) Frequency of Tc1-*mariner* transposon elements in the rainbow trout genome. In B and C, red = high, yellow = medium, green = low sequence similarity or frequency, respectively. D) High-resolution female (red) and male (blue) linkage maps constructed from the analysis 44,910 markers genotyped in a family material of 5,716 fish.

Linkage mapping detailed striking recombination differences between the sexes across the genome (Figure 1d; Supplementary Information section 3), resolved variable chromosome numbers associated with centric fusions or fissions in rainbow trout (Extended Data Figure S2), and revealed recombination interference suggestive of large polymorphic inversions on chromosomes Omy05 and Omy20 (Figure 2; Extended Data Figure S3). Haplotypes tagging these rearrangements were identified and utilized to classify the parents of the mapping families. Subsequent linkage mapping in homozygous parent families disclosed the structure of the inversions, while linkage mapping in families from heterozygous parents documented almost complete repression of recombination across the rearrangements (Extended Data Figure S3). The Omy05 rearrangement is characterized by two adjacent inversions of 22.83 and 32.94 Mb, of which the first is pericentric, reversing the centromere. The alternative karyotypes in the Omy05 double-inversion were categorized as ancestral (*A*) or rearranged (*R*) based on their sequence and structural synteny relative to the Atlantic salmon, coho salmon, and Arctic char genomes, and linkage maps for Chinook, chum, and sockeye salmon (Extended Data Figure S4).

**Figure 2.**
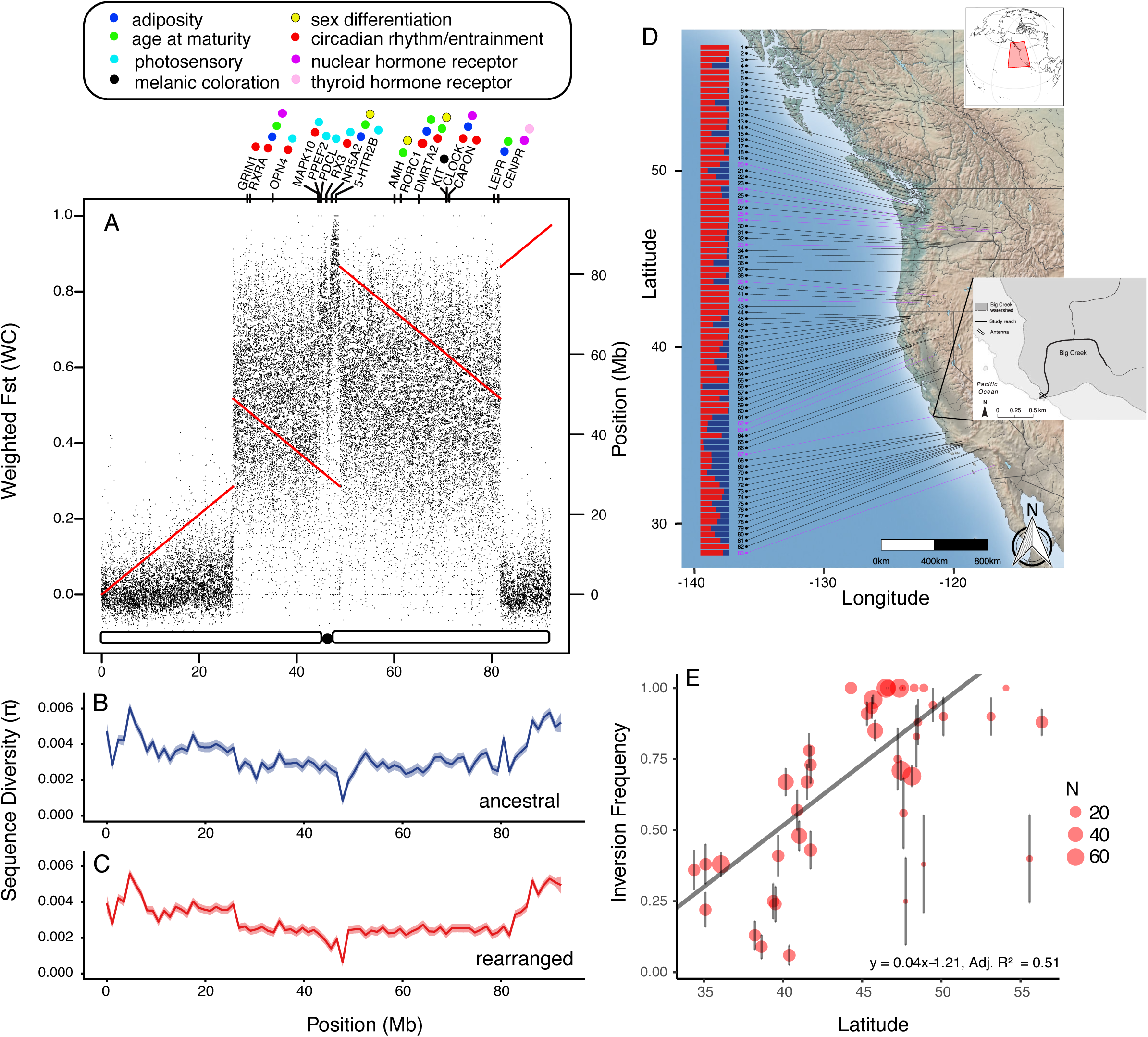
A) Fst across omy05 reveals large area of elevated divergence coinciding with two large linkage discontinuities indicative of two large inversions, the first of which is pericentromeric. Centromere position is illustrated at the base of the plot. Key candidate genes with related functions are spread across the two inversions, colours depict trait relevant functions. B&C) Sequence diversity (π) among AA (blue) and RR (red) individuals, respectively. D) Map of western North America showing all 83 sampling locations, with bars showing frequencies of the A (blue) and R (orange) Omy05 rearrangement karyotypes. Red numbers indicate locations where whole genome resequenced individuals were selected. Map insets: global location and location of Big Creek, Monterey County CA. E) Inversion Frequency as a Function of Latitude among subset of 42 populations of North American rainbow trout with migratory access to the ocean. Point sizes are proportional to sample size with bars showing +/-standard error (SE). Weighted least squares regression line, *y* = 0.04*x* – 1.21, adjusted R^2^ of 0.51.

Whole genome resequencing of 38 individuals across the species’ native range (Table S4) revealed decreased diversity within (Π) and increased divergence between (*FST*) AA and RR karyotype individuals (Figure 2a-c; Supplementary Information section 6, Table S4). A sharp rise in divergence occurred at the inversion boundaries coincident with the break points in the linkage map, with maximal divergence in the pericentromeric region peaking in a 2.5 Mb region of inversion 1 (Figure 2a). This pericentromeric region also displayed a pronounced decrease in sequence diversity (Figure 2b,c) and was enriched for genes containing segregating missense mutations (pericentromeric region 47% versus rest of rearrangement, 30.5%). Dating of coding sequence (CDS) divergence across the inversions suggested that they have been maintained for ~1.5MY (Extended Data Figure S5; see Methods). There was no evidence for different ages of the two inversions, leaving the order of occurrence unresolved. Furthermore, no evidence of differences in the dating estimates was found between the centre and inversion breakpoints either from dating of coding regions or *FST*, consistent with the double inversion forming a very strong barrier to recombination.

## Sexual conflict and sex-dependent dominance

Sex-specific migratory optima are expected to result in intra-locus sexual conflict in rainbow trout where there is a shared genetic basis to migratory tendency. Previous studies have associated genetic markers on Omy05 with migratory traits in rainbow trout^27,28^, and so the double-inversion could potentially have sex-specific effects on life-history. We, therefore, tested the role inversion karyotype plays in mediating sexual conflict and whether sex-dependent dominance contributes to its resolution. Mark-recapture analysis of >2,600 individually-tagged *O. mykiss* from a small stream (Big Creek, California, USA, Figure 2d) showed that sex and Omy05 karyotype (AA, AR, or RR) both significantly influence the probability of an individual migrating to the ocean (Figure 3). These results are consistent with shifts in sex and Omy05 karyotype frequencies within the stream among trout across the migratory size range (>100mm; Figures 3b and 3c). The best-fit model of migratory tendency included sex-dependent dominance of the Omy05 rearrangement, where the karyotype with the highest predicted fitness for each sex is dominant in that sex (ΔAIC 4.68; Supplementary Information section 7, Table S6). At peak size for juvenile marine migration (~150 mm fork length), homozygous AA females were more than twice as likely to be detected emigrating as RR females (70.9% vs. 26.7%), with a reduced difference in males (45.3% vs. 31.8). In the full genotype model, heterozygous females were estimated to emigrate at the same rate as homozygous AA females (complete dominance, d/a=0.97; Figure 3a) while heterozygous males more closely resembled RR males (partial dominance, d/a=0.48; Table S6; Extended Data Figure S6). Such asymmetric sex-dependent dominance is predicted when the strength of antagonistic selection differs between the sexes^9^ as was observed in age at maturity in Atlantic salmon^4^.

**Figure 3.**
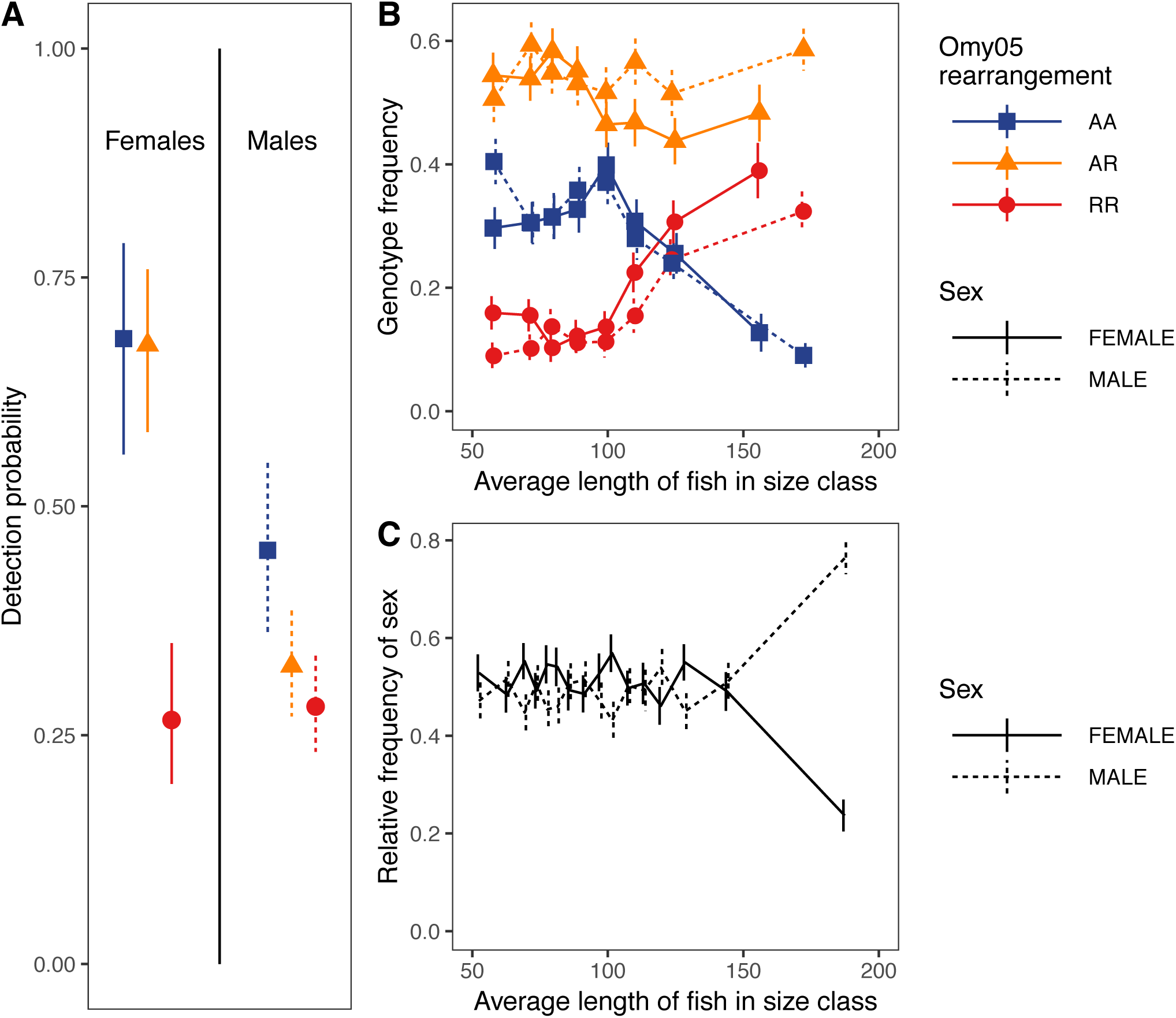
A) Migration of juvenile females (left) and males (right) tagged in freshwater with AA (blue squares), AR (orange triangles), and RR (red circles) genotypes at peak size for individual migration, as estimated from detections at the in-stream fixed antennas, with generalized additive model fits (95% confidence intervals). Proportions of B) Omy05 rearrangement genotypes, and C) Genetic sex, among all females (solid lines) and males (dashed lines) sampled in Big Creek and ordered by size. For B and C, individuals were binned into length categories of ~170 individuals to calculate mean values, with bars showing +/-1 s.e.

The sex-dependent dominance of the Omy05 double-inversion and its role in resolving sexual conflict over migratory tendency could reflect the homeology between the centromeric half of the sex chromosome (Omy29:0-26.28Mb) and Omy05, including 6.45Mb of inversion 1 (Figure 1; Extended Data Figures S4, S7). However, it is unknown what role, if any, the sex chromosomes of rainbow trout play in sexual conflict resolution. The salmonid sex-determining locus, *sdY*^29^, is located within a transposon cassette enabling frequent sex chromosome turnover^21^, evident in among-species sex chromosome transitions^19^, and suggesting the sex chromosomes may not be able to protect sexual conflict polymorphisms like heteromorphic sex chromosomes^5^. To test if homeology with the sex chromosome could explain the role played by the Omy05 supergene in sexual conflict resolution, we next tested for signatures of sexual conflict resolution by the sex chromosomes in rainbow trout.

Elevated genetic differentiation (*FST*) between the sexes is predicted around sexually antagonistic loci in linkage disequilibrium with the sex determiner^3,30^. High between-sex *FST* among 38 wild individuals at Omy29:5Mb as well as BAC sequence alignment supported this region as the location of the *sdY* transposon cassette^31^ (Supplementary Information section 5) and places *sdY* within the region with homeology to Omy05, but outside the rearrangement. Linkage analysis confirmed that male recombination is strongly localized towards the telomere on Omy29 (Figure 4d; Extended Data Figure S7a). Thus, if X-Y divergence is limited only by male recombination, divergence should be seen across most of the Y chromosome. In contrast, divergence was largely restricted to the region between *sdY* and the centromere, where recombination is also low in females (Figure 4a,d). These results suggest that X-Y differentiation is limited over the majority of the chromosome by recombination in sex-reversed males (XY females)^22^, maintaining a large pseudo-autosomal region with limited potential to resolve sexual conflict.

**Figure 4.**
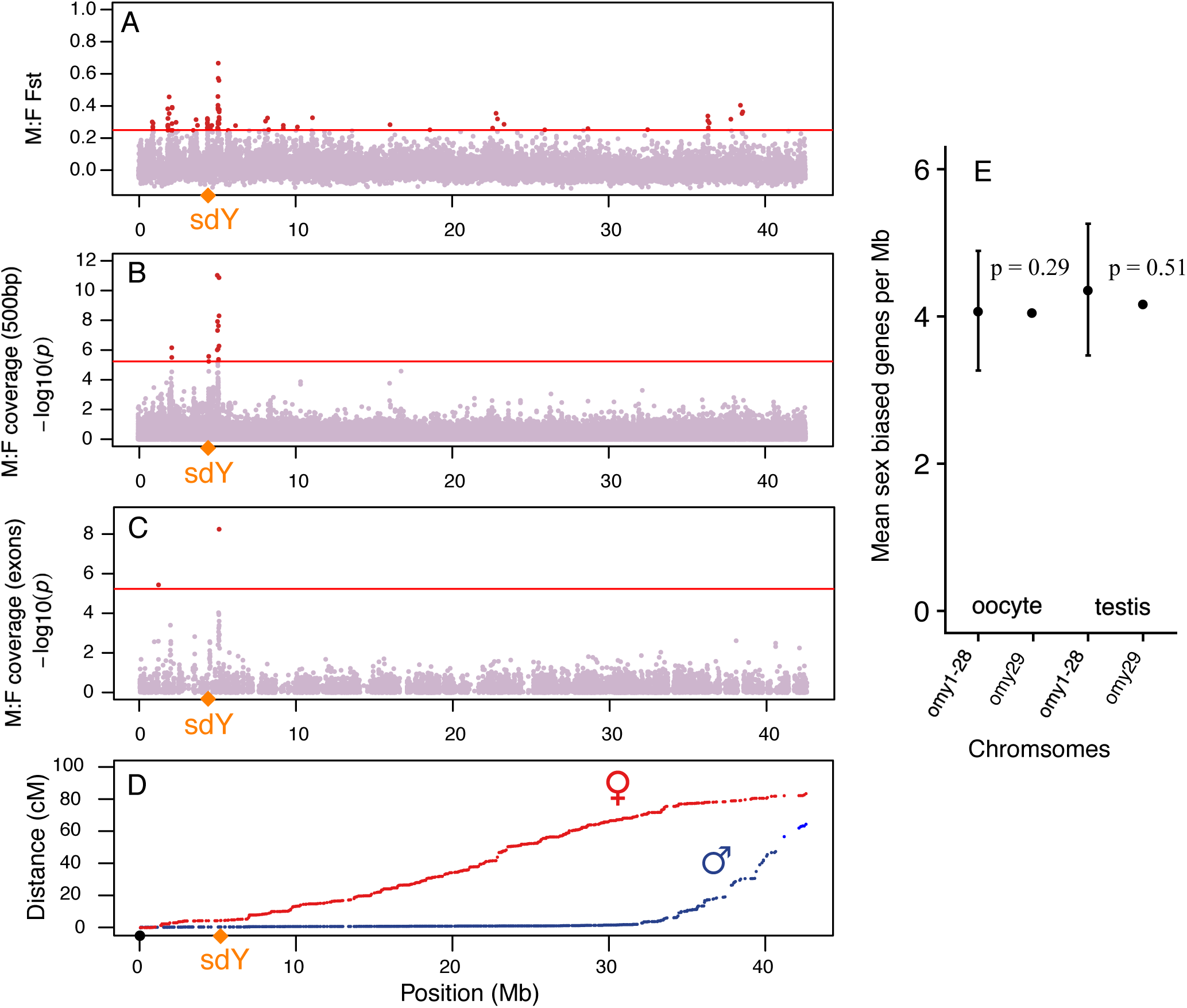
Between sex divergence along the sex chromosome, Omy29, shown by A) Fst, B) coverage in 500bp windows and C) exon coverage is concentrated in the 5Mb region between the centromere (0Mb) and the male determiner, *sdY*. D) This region has low recombination in both males and females. E) Relative to the autosomes, genes with maximal expression in oocyte or testis are not enriched on the Y chromosome.

Y chromosomes typically become enriched with male-specific genes through preferential retention and translocation^32^. However, gene content differences reflecting Y enrichment were not evident in mean coverage between male and female reads across the Y. Instead, coverage differences were only found close to *sdY* (Figure 4b,c), consistent with the Y having not gained or retained genes not present on the X. Two exons had coverage differences between males and females, both in tRNAs which are enriched close to *sdY*, and so may reflect assembly collapse (Figure 4c). To test for Y-chromosome gene loss, we compared its sequence with the orthologous autosome in coho salmon, Oki29^33^. Gene losses between Oki29 and Omy29 were nearly symmetrical (15% and 12% respectively), indicating that gene conservation on the rainbow trout Y chromosome is approximately that expected for an equivalent autosome undergoing rediploidisation. Together these comparisons suggest an absence of the preferential gene gain/retention or loss from the Y chromosome expected under the classical model of sex-chromosome differentiation.

To further test for enrichment of genes with sex-specific gene effects, we defined male and female benefit genes as those with maximum expression in testis and oocytes, respectively, compared to a panel of 13 somatic tissues. The Y chromosome was not enriched for male benefit genes (178/919 Y genes compared to 9005/47415 autosomal genes, -log2 transcripts per million (TPM), one-tailed Fisher’s exact test p = 0.513, Figure 4e), consistent with X-Y recombination and/or frequent turnover preventing enrichment of male benefit genes. Conversely, low recombination between X and Y with weak sexual antagonism can lead to feminisation of the Y chromosome^8^. However, there was also no enrichment in female benefit genes (173/919 Y genes compared to 8091/47415 autosomal genes, -log2 TPM, Fisher’s exact test p = 0.29), with the same proportion of both testis and oocyte genes as expected if they were evenly distributed across the chromosomes (18-19 %, Figure 4e). These results show a lack of gene content bias towards either male or female benefit genes, in contrast to previous findings^32,34^, suggesting that the Y chromosome has not accumulated either male or female benefit genes.

Finally, we found little evidence of structural rearrangements on Omy29 that are predicted to accompany Y chromosome recombination shutdown^35^. Instead, Omy29 chromosome structure is highly conserved with the distantly related Atlantic salmon and charr (divergence times ~20 MYA; Extended Data Figure S7c,d), whereas the comparative regions in all other mapped Pacific salmon species are rearranged (Extended Data Figure S7e,f). Within rainbow trout the homeologous region on Omy05 is highly rearranged, creating a barrier to homeolog recombination with Omy29, particularly in the region of *sdY* (Extended Data Figures S4a,c; S7b). This difference between homeologs likely predates the speciation of salmonids as collinearity is maintained between Omy05 and its homologs in Atlantic salmon and charr (Extended Data Figure S7).

The absence of XY differentiation, sex-biased gene enrichment, rearrangements, or other signals predicted by the classical model of sex chromosome evolution^18,22^ suggests that selection against deleterious mutation load rather than the resolution of sexual conflict has driven Y chromosome evolution in rainbow trout. In contrast, the absence of homozygote lethality of either karyotype of the Omy05 inversion allows within-karyotype recombination to purge deleterious mutations, avoiding degradation. Thus, sex-dependent dominance of the Omy05 rearrangement provides an alternative, autosomal mechanism of chromosomal scale sexual conflict resolution that avoids the mutation load associated with the canonical heteromorphic sex chromosome system.

## Gene composition of the Omy05 rearrangement

Inversion supergenes allow coadapted variants to avoid being broken up by recombination^36^. The 56Mb double-inversion contains 1,091 protein coding genes, including key adiposity, circadian rhythm/entrainment, photosensory, and age at maturity genes associated with migratory behaviour and seasonal timing of maturation in salmonids (Figure 2a, Table S7; see Methods). These include the master regulator of circadian rhythm, *CLOCK*, and a visual pigment, OPN4, expressed in the saccus vasculosus, the organ that controls photoperiodism and so reproductive timing in fish^37^. Additionally, multiple sex determination/differentiation genes are found, notably a doublesex/male abnormal 3 (DM) domain-containing transcription factor, *DMRTA2*.

*DMRTA2* is a vertebrate duplicate of *Doublesex*, a gene known to be responsible for sexual dimorphism in insects including an autosomal sex-limited butterfly mimicry supergene^38^. *DMRTA2* is expressed in both the gonads and developing pituitary of zebrafish, regulating terminal differentiation of corticotropes and gonadotropes, and may accelerate gonadal development through increases in luteinising hormone and follicle stimulating hormone and so influence maturation timing^39^. Interestingly, *DMRTA2* morphants exhibit reduced numbers of cells expressing pro-opiomelanocortin (*pomc^39^*), which is differentially expressed between freshwater migratory marine forms of rainbow trout^40^ and Atlantic salmon^41^. Two other genes known to strongly affect sex-specific development are found in the inversion, *AMH*^42^ and *NR5A2*^43^. All three of these genes also influence maturation (*DMRTA2*, *AMH*^39,44^ and age at maturity (*NR5A2*^45^).

Photosensory function is critical for seasonal timing of smoltification and maturation, and visual-related pathways are over-represented in gene expression differences between resident and anadromous rainbow trout^40^. The Omy05 rearrangement harbours multiple genes related to photoperiodic circadian entrainment, including a key non-image forming photosensory gene linked to seasonal timing, *OPN4*, known to be expressed in the coronet cells of the saccus vasculosus^37^, and the photic circadian entrainment genes *GRIN1*(*NMDAR*)^46^ and *CAPON^47^*. Additionally, the highly divergent pericentromeric region of the Omy05 rearrangement contains a cluster of genes with both photosensory and circadian functions, *PHLP*, *PPEF2*, *RX3,* and *MAPK10*^48-50^ (Figure 2a)*. Rx3* mutant zebrafish display severe attenuation of the cell cycle rhythms and an absence of *pomc-*expressing cells^49^. A missense mutation within *RX3* was predicted to have an effect on phenotype (polyphen2 analysis Thr279Ile, naïve Bayes posterior probability 0.846, sensitivity = 0.83, specificity = 0.93). Additionally, the serotonin receptor, *5-HT2B*, a neurotransmitter with effects on both behaviour and retinal development^51,52^, contains two segregating missense mutations. The homeolog of the pericentromeric region on Omy12 has also been implicated in migratory phenotype^53,54^, supporting a role for these genes and that differences in both the phenotype and visual signalling can also be generated independently of the Omy05 rearrangement via parallel selection on gene duplicates. Several major QTL for migratory traits are located on the homeologs of the Omy05 rearrangement in Northern populations (e.g. *CLOCK* (Omy01)*, MAPK10* (Omy12); Nichols *et al*. 2008; Hecht *et al* 2013) suggesting a potential shift to a more diffuse architecture in the north.

The intricate feedbacks between adipogenesis and circadian rhythm are increasingly recognised^55^, and adiposity is consistently associated with divergent migratory phenotypes in rainbow trout and age at maturity in fish^4,10,56^. Multiple genes within the Omy05 rearrangement are major adiposity-related genes, e.g. *RORC1, RXRA, NR5A2, LEPR*, all of which are also associated with age at maturity in humans^45^*. RORC1* plays a central role in the interactions between lipid metabolism, homeostasis, and circadian rhythms^55^. Similarly, *CLOCK* regulates adipogenesis^57^ and contains two missense mutations that perfectly segregate with the *A* and *R* haplotypes, one of which is predicted to affect the phenotype through loss of a glutamine residue in the polyQ tail (polyphen2, ancestral to rearranged: His871Gln (Gln>His) naïve Bayes posterior probability 0.641, sensitivity = 0.87, specificity = 0.91). *NR5A2* is involved in oestrogen biosynthesis (Chiang *et al* 2000) and inhibition of adipogenesis through regulation of *CYP19a1a*, expression of which falls when that of the master regulator of adipogenesis, *PPAR*-gamma, increases^58^.

Prior to migration, individual rainbow trout transition from a riverine melanotic to a marine cryptic silver coloration. KIT signalling is required for the establishment and survival of melanophore progenitors^59^, and KIT is located adjacent to CLOCK in the inversion (Figure 2a). Finally, thyroid hormone signaling is known to control both maturation and smoltification in salmonids, and *CENPR,* a coactivator of oestrogen receptor alpha^60^ and enhancer of nuclear receptors, thyroid hormones, and retinoid X receptors (*RXR*^61^), is found near the distal break point of inversion 2 (Figure 2a), providing an important link between sex steroid control, adiposity, and circadian rhythms.

Within such a large double-inversion it is unlikely that a single causative gene or mutation is responsible for the karyotype:phenotype association with the many fixed differences making it challenging to identify specific causative elements. However, *DMRTA2* may play a critical role in the sex-specific effects of the double-inversion, as a the control of sex-limited traits is a key function of DMRT transcription factors that is conserved across the animal kingdom^39^.

### Geographic distribution of the Omy05 rearrangement

The frequency of the rearranged karyotype varied with migratory access among 91 geographically distributed populations (mean R frequency above vs. below barriers, 0.84 vs. 0.66, p < 0.001; Figure 2d; Table S4; Extended Data Figure S8), as previously observed in the south^28^, confirming that the link with the migratory trait extends across the species range. However, among the 42 ocean-accessible populations, we observed a strong cline in the R karyotype with both latitude (adjusted R^2^ = 0.48, p < 0.001; Figure 2e) and monthly mean ambient temperature (Max. adjusted R^2^ = 0.61, p < 0.001; Extended Data Figure S9). These patterns are likely driven by temperature dependent developmental rates^10,62^, major QTL for which overlap the Omy05 rearrangement^27,54,62^, strongly predicting reproductive timing and early male sexual maturation^63^. Thus, despite the strong influence of the Omy05 double-inversion on individual migratory tendency in Big Creek, near the southern extent of the species’ range, this effect is likely reduced where the ancestral, slow development, karyotype is temperature-limited.

In these colder, high latitude populations, the faster intrinsic development rate required to compensate for the effects of decreased temperature may result in positive selection for the rearranged karyotype irrespective of migratory phenotype (Extended Data Figure S10). However, anadromy also becomes rarer at high latitudes, reflecting both the increased food availability in rivers with large salmon runs and the increased accumulation of adipose tissue with lower metabolic rates in cooler rivers^10,64^. These trends reduce the cost of residency for females, suggesting a concomitant decrease in sexual antagonism and that sexual conflict resolution by sex-dependent dominance is dependent on geographical variation in the strength of sex-specific selection^1,4,65^.

### Insights

Inversion supergenes maintained by balancing selection have been associated with the evolution of novel dominance patterns, and unlinked modifiers are thought to act epistatically on inversion effects^16,36,38^. Our results demonstrate, for the first time, sex-dependent dominance of an autosomal inversion contributing to the resolution of sexual conflict over a complex life-history trade-off. Sex-dependent dominance of the higher fitness karyotype within each sex provides the conditions for net heterozygote superiority^7^, allowing independent optimisation of the migratory phenotype in both males and females and the maintenance of sexual conflict polymorphisms over a large chromosomal segment. In stark contrast, the Y chromosome lacks signatures typical of sex chromosome evolution, with no evidence for enrichment of male benefit genes, isolating structural rearrangements, or an excess of gene loss indicative of degeneracy. These patterns strongly suggest that Y chromosome evolution in rainbow trout is driven primarily by the avoidance of mutation load, with limited capacity to maintain sexual conflict polymorphisms and that the Omy05 rearrangement represents an autosomal alternative to the canonical model of sexual conflict resolution by sex chromosomes.

Linkage is expected to both develop among sexually antagonistic loci and to relax the conditions for maintenance of sexually antagonistic variation because of increased fitness effects^66^. Such processes can lead to the accumulation of loci with large fitness effects predicted under sexual antagonism^67^, and so the sex-dependent dominance of an autosomal supergene we observe in rainbow trout may represent a common mechanism for the maintenance of polygenic sexually antagonistic variation.

In contrast to other inversions underlying alternative reproductive tactics^12-15^, lack of homozygous lethality enables the Omy05 rearrangement to purge deleterious mutations from both karyotypes^8^. The inversion complex has been maintained by sexually antagonistic balancing selection for ~1.5 MY, and so represents a stable alternative mechanism of sexual conflict resolution that avoids costs of mutation load accumulation by sex chromosomes. However, this architecture is likely geographically restricted by counter-gradient selection on temperature-dependent developmental rate and changes in the strength of sexual antagonism with latitude. The maintenance of fitness variation by sexually antagonistic selection has important conservation implications, however, geographical variation in the strength of such selection highlights the complexity involved in incorporating adaptive genomic variation into conservation management^68^.

## Supporting information

Pearse_etal_SI

Pearse_etal_Additional_Figures

Pearse_etal_TableS3

Pearse_etal_TableS4

Pearse_etal_TableS7

## Methods

### Genome sequencing and assembly

We sequenced and assembled the rainbow trout genome using DNA from a single homozygous doubled haploid YY male from the Swanson River (Alaska) clonal line (BioProject <u>PRJNA335610</u>) using a complementary combination of inputs, including short-read sequencing technology from Illumina, high-density linkage mapping, and the DeNovoMAGIC genome assembly pipeline from NRGene^69^, as well as long-range data from Dovetail Genomics Chicago library sequencing^70^ and comparative genomic information from the Atlantic salmon genome^24^ (Supplemental Information section 1.1). To anchor, order, and orientate scaffolds into chromosome sequences, we constructed a high-density linkage map using Lep-MAP software^71^ from a pedigree of 5,716 fish genotyped with a 57K SNP-array^72^.

Repeat masked chromosome sequences for rainbow trout were aligned against each other using LASTZ^73^ to identify 98 homeologous blocks originating from the Ss4R (for details see Supplementary Information section 2). Sequence similarity between homeologous sequences was determined in 1□Mb intervals by averaging local percentage of nucleotide sequence identity using high-scoring segment pair (HSP) from LASTZ alignments and presented as a Circos plot^74^ in Figure 1.

### Whole genome re-sequencing

Genomic DNA was extracted from fin clips of 61 rainbow trout. Whole-genome paired-end sequencing libraries were prepared and sequenced using the Illumina HiSeq 2000 and 2500 platforms providing an average of 15X and minimum of 8X genome coverage per sample. The sequenced samples included 11 clonal lines from Washington State University, 38 steelhead and resident rainbow trout from wild and hatchery-origin populations distributed throughout the native range of the species, and 12 fish from the AquaGen rainbow trout aquaculture breeding program. The origin and geographic location of each sample are given in Supplemental Table S4. The bioinformatic pipeline for mapping of the resequencing data to the reference genome, variant calling, and quality filtering of the SNPs was previously described^75^. A total of 31,441,105 SNPs were identified genome-wide, of-which the subset of SNPs that mapped to chromosome Omy05 was further used for population genetic and functional analyses as described below.

### Orthologs identification

Orthologs were first identified using orthofinder^76^ with protein sequences of two mammalian species (mouse and human) and eight teleost species (zebrafish, medaka, stickleback, northern pike, coho salmon, rainbow trout, European grayling, and Atlantic salmon) as the input. Gene annotations for mouse, human, zebrafish, medaka, and stickleback were downloaded from ENSEMBL. For northern pike and Atlantic salmon we used the NCBI RefSeq annotations for assembly versions ASM72191v2 and ICSASG_v2, respectively. For coho salmon we used TransDecoder (https://github.com/TransDecoder/TransDecoder/wiki) to predict protein sequences based on a *de novo* transcriptome assembly. Rainbow trout gene annotations were based on an in-house annotation pipeline previously described and used for Atlantic salmon^24^.

For each orthogroup, protein sequences were aligned using MAFFT and protein trees were estimated using FastTree. Orthogroup protein trees containing duplication nodes ancestral to all teleosts or vertebrates were then further partitioned into smaller clan-trees using an in-house R-function (available from gitlab: https://gitlab.com/sandve-lab/salmonid_synteny/blob/master/clanfinder_function.R). New codon alignments for orthogroups, including those converted to ortho-clans with reduced gene tree complexity, were made using Pal2Nal to convert protein alignments into CDS alignments. These nucleotide alignments were then used to re-estimate a final set of ortholog gene trees.

### Generation of time-calibrated Beast gene tree for genes on Omy05

For each gene on Omy05 we identified the corresponding gene tree and performed a series of filtering steps to retain only orthogroups for which we had high confidence classification of both orthologs and duplicates originating from the salmonid whole genome duplication. This was ensured by filtering according to gene tree topologies using the following criteria: (i) salmonid tips had to be monophyletic, (ii) gene tree phylogenies had to conform to the species phylogeny for non-salmonid taxa (after rooting in the most distant salmonid outgroup), (iii) gene tree had to have retained all duplicates from the salmonid whole genome duplication (Ss4R), (iv) the topology of Ss4R duplicates had to conform to the salmonid species phylogeny.

After identifying high confidence gene ortholog groups for rainbow trout genes on Omy05 we made sequence alignments for these ortholog groups using both CDS and whole genes containing introns. Coding regions (CDS and genes) were extracted from the whole genome sequence vcf file using vcftools and then refilled to sequence using the vcf2fasta function in FreeBayes^77^ using gene fasta files created with samtools^78^. Prior to alignment we swapped out the single rainbow trout nucleotide sequence from Omy05 with two sequences corresponding to the two inversion haplotype sequences. Sequences from each orthogroup were then aligned with MAFFT using default parameters and alignments were trimmed using gblocks^79^.

Finally, Beast^80^ was run on each alignment (CDS and gene) which had >0 base pair differences between the inversion haplotypes using the parameters chainLength=10000000, storeEvery=5000, a Yule-model of speciation, the HKY substitution model and a relaxed molecular clock. For priors we assumed: 1) the duplicate salmonid gene lineage including the Omy05 haplotypes to be monophyletic and with a divergence time of 20 million years ago (log-normal distribution with standard deviation of 0.1); and 2) northern pike and all salmonids to be monophyletic and diverging 125 million years ago (log-normal distribution with standard deviation of 0.1).

### Sex chromosome evolution

#### Between sex divergence (*F_ST_*)

Between sex Weir and Cockerham’s *F_ST_* was calculated for each SNP from the whole genome resequence dataset and also separately for wild individuals, pooled across populations, using *vcftools*. The inclusion of YY double haploid males increased the peak height in the sequence surrounding the *sdY* location, but this was also the highest between-sex *F_ST_* peak in the wild individuals. Between-sex *F_ST_* was plotted for Omy29 and compared with the linkage map for this chromosome (see above).

#### Coverage

Genotypic sex was determined by coverage of the *sdY*-containing scaffold, *KJ851798.1:20860-23610,* where samples with a coverage of zero were typed as female. Samples were processed using the Speedseq pipeline^81^ and aligned with BWA-MEM^82^ version 0.7.10-r789. Coverage data was calculated using Mosdepth^83^ version 0.2.1, both for the whole genome in 500 bp windows and for all exons from the RefSeq100 Omyk_1.0 annotation. For Mosdepth, the mapping quality threshold was set to 10. Log_2_ coverage was plotted and the significance of the difference in coverage was calculated by a t-test.

### Expression (RNAseq)

RNAseq data from 15 tissues (spleen, kidney, gill, head kidney, skin, intestine, liver, red muscle, white muscle, brain, fat, stomach, pineal, oocyte and testis) were downloaded from SRA. Quantification of RNA-seq data was performed using Kallisto^84^ version 0.44.0, with 30 bootstrap samples. We defined testis and oocyte genes as those expressed maximally in testis and oocytes, respectively, and had a TPM >5. Enrichment of these sex-specific genes on the Y (Omy29) was tested with Fisher’s exact tests.

### Gene gain and loss

The gene content of the orthologous rainbow trout Omy29 and coho Oki29 chromosomes were compared using a combination of gene orthology relationships determined by orthofinder (https://doi.org/10.1186/s13059-015-0721-2), see ortholog relationships above, and protein BLAST. We used BLAST to increase the number of genes with defined ortholog relationships between the two chromosomes. Where the matching gene mapped to an unmapped scaffold and the duplicate mapped to the homeolog of chromosome 29 in the other species, we assumed the unmapped copy belonged on chr29; this may mean we miss some translocations away from chr29, but since there were more unmapped chr29 genes in coho than in rainbow trout we considered it conservative for understanding gene content evolution in rainbow trout.

### Genome Sequence Synteny

Rediploidisation of salmonid genomes has progressed through the rearrangement and fission/fusion of large chromosomal blocks with identifiable syntenic relationships of these blocks among species^24^. Large scale chromosomal synteny was determined through the comparison of published RADseq linkage maps for coho^85^, Chinook^86^, chum^87^, and sockeye salmon^88^, and genome sequences available for Arctic char (GCF_002910315.2), coho (GCF_002021735.1) and Atlantic salmon (GCF_000233375.1).

### Field sampling and capture-recapture experiment

#### Fieldwork

The study was conducted in a natural population of *O. mykiss* in Big Creek, a small (58 km^2^) coastal watershed along the central California Coast in Monterey County, California, USA.

Individuals in this population have free access to migrate to the ocean, but many mature residents remain in the creek^89^. We non-lethally captured, weighed, measured, and took caudal fin samples from more than 2,600 individuals in ~1,900 m of contiguous stream habitat starting at the Pacific Ocean entry of Big Creek between May 2006 and Oct 2009. All fish >100 mm fork length (FL) were injected with unique passive integrated transponder (PIT) tags prior to release at the capture location. Tagged fish were available for detection by a continuously active in-stream fixed antenna ~70 m upstream from the ocean that recorded the date and time of migration by detected individuals between May 2006 and Oct 2012. Due to tag size limitations, only individuals >100mm FL received a tag (23-mm half-duplex) detectable by the antenna. The length and weight assigned to tagged fish detected migrating from the stream were based on measurements taken at their last physical capture, typically the autumn prior to migration. Analysis of antenna detections considered the subset of fish (n = 887) that were 1) tagged with PIT tags detectable by the antenna, 2) captured ≥ 100 m upstream from the antenna array near the stream mouth and, 3) successfully genotyped for Omy05 and OmyY1 sex identification loci. From this group, *migrants* were defined as individuals detected at least once by the in-stream antenna that 1) had no subsequent capture or detection, 2) were at large <730 days between their last capture and final detection, and 3) were detected at the array <15 days in total and over a span of <60 days. These criteria were used to distinguish detections of true migrants (i.e. smolts, N = 331) from detections of tags no longer in live fish (e.g., following death or predation of a tagged fish; N = 68). The remaining 488 individuals were never detected by the in-stream fixed antenna, so are assumed to be either non-migratory residents or in-stream mortalities.

### SNP Genotyping

Genomic DNA was extracted from fin clips of all fish sampled in Big Creek and genotyped at 95 SNPs^90^ using TaqMan assays (Applied Biosystems, Inc.) on 96.96 Dynamic SNP Genotyping Arrays with the EP1 Genotyping System (Fluidigm Corp.). An additional TaqMan assay was designed around a Y chromosome-linked gene probe^91^ and an invariant autosomal control gene and was used to determine genetic sex. Two negative (no DNA) controls were included in each array, and genotypes were called using SNP Genotyping Analysis Software v3.1.1 (Fluidigm Corp.). Two loci located within the Omy05 rearrangment, Omy_114448-87 and Omy_121006-131, show strong linkage disequilibrium with each other and with other loci located within the Omy05 rearrangement^28^. In Big Creek, these two loci are in near-perfect disequilibrium, with 97.2% identical genotypes. We designated the C and T bases at SNP Omy_114448-87 as *A* (ancestral / anadromous) and *R* (rearranged / resident), respectively for the Omy05 rearrangement. The remaining 92 non-Omy05 loci were used for population genetic and kinship analyses.

### Capture-Recapture Modelling

The effects of Omy05 inversion haplotype on migratory behavior were investigated using a channel-spanning PIT tag in-stream antenna ~70 m upstream from the ocean. We predicted that the probability of detecting anadromous tagged fish would be highest for fish tagged near the size that migratory steelhead smolts typically emigrate from freshwater^92^ (~150 mm FL). Detection probability should be lower for anadromous fish sampled at a smaller size, since many will not survive to emigrate, and also for larger fish, since they are presumably maturing to remain as residents. Fish with genotypes favoring residency may also be observed at the antenna, but we expect this probability to show no peak near 150 mm FL. To test these hypotheses, we used generalized additive models to estimate the probability of detection at the downstream antenna as a function of length at last capture and release, with Omy05 genotype and sex as categorical covariates, using the mgvc package (version 1.8-4) in the statistical programming language R (version 3.1.3). Because earlier work showed that the sex ratio of fish smaller than 150 mm was near 50:50 while larger size classes are enriched in males^89^, we expected that there should be an interaction between sex and genotype. We therefore tested seven models capable of accommodating increasing variability. The simplest model assumed emigration probability is only a smooth function of length at tagging and release. Slightly more complex models included effects of sex and genotype separately (two and three smooth functions, respectively). The final four models treated the sex and genotype interaction differently, allowing for partial dominance (six smooths, one for each sex and genotype combination), anadromous or resident dominance, or sex-dependent dominance reversal, with anadromy dominant for females and residency dominant for males. The best model according to Akaike’s Information Criteria included sex-dependent dominance reversal, with an AIC 4.68 lower than the next highest-ranked model (Table S6).

### Geographic survey of Omy05 variation

#### Omy05 SNP populations survey

Tissue samples (~1-3 mm^2^) or genomic DNA were obtained from 63 natural origin *O. mykiss* populations (*N* = 1,592 individuals) ranging from the Kamchatka Peninsula, Russia to southern California, USA. Also, 94 samples from four steelhead hatchery broodstock and 141 samples from six rainbow trout hatchery strains were genotyped (Supplemental Table S4). These samples were all genotyped using a separate panel of 86 SNPs distributed on chromosome Omy05—55 SNP assays from Pearse *et al*.^28^ and 31 developed from Miller *et al.*^62^. Eight loci were outside the inversion region and three failed to amplify, leaving 75 SNPs retained for analyses. Allele frequencies were assessed using the R package *adegenet* 1.3–4^93^. Linkage disequilibrium was estimated using the allelic correlation coefficient r-squared^94,95^ using the R-based package *genetics*^96^. The number of pairwise linkage associations that were above a critical value of 0.95 were counted for each population. Because pairwise r-squared values cannot be estimated in populations where either locus is monomorphic, we divided the number of values above the critical value by the total number of r-squared values calculated for each population to obtain a weighted value. We used level plots in R to visualize the allele frequency and linkage disequilibrium data.

Of the 1,592 individuals examined, 286 fish from 33 sampling locations were homozygous AA, and 846 fish from 57 sampling locations were homozygous RR. Population level trees were generated with *poppr*^97^ v. 2.3.0 in R (v. 3.3.1) with Rstudio (v. 1.0.136) with the *aboot* command. Chord distances between populations^98^ were calculated and a tree constructed with the Neighbor-Joining algorithm^99^. Confidence of inferred relationships was evaluated with 1,000 bootstrap replicates.

Finally, the population SNP survey dataset was combined with additional data on inversion frequencies for a combined total of 2,249 individuals from 91 populations (Supplemental Table S4). For all data sources, inversion frequency was inferred based on common SNPs that perfectly or nearly-perfectly identified inversion haplotype.

#### Inversion Frequency as a Function of Latitude

A plot of latitude (*x*–axis, WGS84) and inversion frequency (*y*–axis) was generated by filtering inversion frequency data for those samples from North America taken below migration barriers and without known hatchery introgression (Figure 2; Supplemental Table S4). Graphics and data analysis occurred in R v. 3.3.1 in Rstudio v. 1.0.136 with the *ggplot2* package v. 2.2.1. For each estimate of inversion frequency, standard error (SE) was calculated with *p* equal to the frequency of the inversion in the sampled population and *N* equals the number of individuals sampled from that population. To the plot of latitude and inversion frequency, a weighted least squares regression line was added from a model with inversion frequency dependent on latitude of the sampled population weighted by sample size of the population. The resulting model (*Inversion Frequency* = 0.04 *x* (*Latitude*) – 1.21) has an adjusted R^2^ of 0.51.

#### Inversion Frequency as a Function of Temperature

Temperature data were obtained for each month of the year from the WorldClim^100^ dataset v. 1.4. The dataset represents current temperatures, taken from interpolations of observed data from 1960 to 1990 within 30 arc-second squares. Temperature data were extracted with the *raster* package v. 2.5-8 in *R.* A bilinear interpolation was implemented to average between the four nearest 30 arc-second squares to the collection location. As with the inversion frequency as a function of latitude, a weighted least squares regression weighted by sample size was conducted for each month of the year and the resulting adjusted R^2^ was retained for each sub-plot (Extended Data Figure S9).

## Acknowledgements

We thank H. Fish, K. Pipal, and many others for fieldwork, V. Apkenas, A. Carlo, and E. Campbell for assistance with data collection and analysis, and M. Readdie and F. Aryas for support at the University of California Landels-Hill Big Creek Reserve. Samples and data for the geographic survey were provided by M. Ackerman, S. Lewis, S. Narum, K. Nichols, S. Northrup (FFSBC), E. Taylor (UBC), D. Teel, and K. Warheit.

Compute Canada provided computing resources used in repeat annotation and analysis. We thank R. Long and K. Shewbridge for their help in DNA sample preparation for sequencing and genotyping and in the preparation of RAD sequencing libraries, and K. Martin and Troutlodge, Inc. for the permission to use samples from their germplasm for genotyping.

We also thank the Genomics Core at Washington State University, Spokane, WA, the University of Idaho Genomics Core and the Vincent J. Coates Genomics Sequencing Laboratory at UC Berkeley for performing DNA library preparation and clonal lines’ re-sequencing. The genome re-sequencing of the Whale-Rock female clonal line was conducted in collaboration with M. Garvin, Oregon State University. Mention of trade names or commercial products in this publication is solely for the purpose of providing specific information and does not imply recommendation or endorsement by the U.S. Department of Agriculture. USDA is an equal opportunity provider and employer. This project was supported by funds from the USDA Agricultural Research Service in-house project numbers 1930-31000-009 and 8082-31000-012. DH clonal line re-sequencing was supported by the Agriculture and Food Research Initiative Competitive Grant no 2015-07185 from the USDA National Institute of Food and Agriculture and by an NRSP8 Aquaculture Genome funding seed grant to M. Garvin and G. Thorgaard. The whole-genome re-sequencing data provided by K. Naish was obtained from a project supported by the Agriculture and Food Research Initiative Competitive Grant no 2012-67015-19960 from the USDA National Institute of Food and Agriculture. Funding for bioinformatics and statistical support at CIGENE-NMBU was provided by NFR grants 208481, 226266 and 275310. Bioinformatic analyses were performed using resources at the Orion Computing Cluster at CIGENE-NMBU, with storage resources provided by the Norwegian National Infrastructure for Research Data (NIRD, project NS9055K). We acknowledge the help of S. Karoliussen and M. Arnyasi at CIGENE-NMBU for generating rainbow trout genotypes and M. Baranski for work on the genetic linkage maps. C. Primmer and K. Nichols provided valuable comments on the draft manuscript.

## Author contributions statement

S.L., Y.P. and A.H. co-conceived the genome assembly project and G.T. provided the Swanson clonal line for the reference genome. Y.P. and T.M. contributed SNP chip and RAD SNP genotype data for the linkage analysis, S.L. and T.M. performed linkage analyses, and T.N. and S.L. refined the assembly and built chromosome sequences. K.B., G.B-Z., D.S-T. and O.B. designed and conducted the DeNovo MAGIC genome assembly of the Swanson clonal line Illumina sequence data. M.M. and L.C. provided RAD SNP data and linkage information for chromosome anchoring of the assembly scaffolds and contigs. D.P. and J.G. contributed the Dovetail sequence data for the genome assembly, and G.G. incorporated the Dovetail sequence data for bridging and combing genome assembly scaffolds. S.L.,Y.P., G.T., B.K., N.B. and D.P. designed the whole-genome re-sequencing study. G.T., B.K., K.K., K.N. M.B., and T.M. contributed samples, data, and/or analysis to resequencing study. D.M. and B.K. created and annotated repeat library and performed the Tc1-Mariner analysis. G.G. performed bioinformatics analyses on the SNP chip genotype and RAD sequence data, and N.B., G.G., and M.C. analyzed the whole-genome resequencing data. S.L. produced data and completed comparative genomic analyses. N.B., M.K., T.N. and S.L. produced and analysed genotype data. N.B. and M.C. performed population genomic analysis of the inversions using resequence data and analysed gene content of the inversions. N.B. and T.N. performed analysis of sex chromosome evolution. S.S., M.K. and T.N. generated RNA data and S.R.S. generated orthogroup gene trees. N.B. and S.S dated the inversions on Omy05. D.R., T.W., D.P., E.A., J.G., and S.T.L. conceived, designed, and conducted the Omy05 capture-recapture field experiment, and D.R. E.A., D.P., and S.T.L analysed the data. A.A-C, J.G., and D.P. conceived, designed, and conducted the SNP populations survey, E.R. and B.K. contributed additional data, and A.A-C., E.A., and M.C. analysed the data. N.B., S.L., E.A, M.C, S.T.L, D.P, and B.K. created figures. D.P., N.B., Y.P., and S.L. wrote the paper with input from all authors. All authors read, commented on, and approve the manuscript.

## Competing interest statement

The authors declare no competing interests. **Animal use**: All animal handling was conducted in accordance with approved institutional guidelines.

