## Supplementary material for "Sex-dependent dominance maintains migration supergene in rainbow trout": Pearse_etal_SI

**Outline:**

1. High-density linkage maps and genome assembly
   1.1. *Sequencing*
   1.2. *De novo draft assembly*
   1.3. *High-density linkage maps*
   1.4. *Construction of chromosome sequences*
2. Post-Ss4R rediploidization characteristics
3. High-density linkage mapping to resolve genome organization
4. Rainbow trout repeats
5. sdY mapping
6. Omy5 re-arrangements and diversity analysis
   6.1. *Detection of haplotypes associated with the two inversions.*
   6.2. *Re-sequencing for SNP discovery and functional analysis.*
7. Fish movement experiment in Big Creek.
   7.1. *Field sampling and capture-recapture experiment*
   7.2. *Population, family, and fecundity effects*

**1. High-density linkage maps and genome assembly**

*1.1. Sequencing*

DNA from the Swanson doubled haploid (DH) YY male line was used as the template for DNA sequencing (BioSample: SAMN05449231). The same DH line was used for the previous genome sequence assembly (GCA_900005705.1; (Berthelot et al. 2014), and for the BAC physical map for rainbow trout (Palti et al. 2009). Two genome shotgun libraries and three mate-pair libraries were prepared for Illumina sequencing. The 450bp and 800bp shotgun libraries were prepared with the Hyper Kapa Library Preparation kit (Kapa Biosystems). The three mate-pair libraries were constructed with the Nextera Mate Pair library Sample Prep kit (Illumina), followed by the TruSeq DNA Sample Prep kit. The 450bp shotgun library was quantitated by qPCR and sequenced on four lanes on a HiSeq2500 for 261 cycles from each end of the fragments using a Rapid TruSeq SBS kit version 2. The 800bp shotgun library and the three mate-pair libraries were also quantitated by qPCR and then pooled and sequenced on four lanes for 161 cycles on a HiSeq2500 using a TruSeq SBS sequencing chemistry version 4. Fastq files were generated and de-multiplexed with the bcl2fastq v1.8.4 Conversion Software (Illumina).

The total number of sequence generated was ~585 billion basepairs providing a hypothetical genome coverage of ~244X. The sequence data is summarized in Supplementary Table S1. Sequence reads were submitted to the NCBI SRA database (Bio-Project PRJNA335610; Study Accession SRP086605; Sample Accession SRS1672116).

***Supplementary Table S1.*** *Sequencing of the rainbow trout genome. The genome of the Swanson doubled haploid (DH) YY male line was sequenced on the Illumina HiSeq2500 using two Paired-end and three Mate-pairs libraries to ~244X genome coverage. The predicted genome size of 2.4 Gb was used to calculated estimated coverage.*

| Library | Type | Insert size | Read length | No. of Paired Reads | No. of Bases | Coverage (X) |
| --- | --- | --- | --- | --- | --- | --- |
| 450bp | Paired-end | 350 - 500 | 2 x 260 | 510,844,344 | 265,639,058,880 | 110.7 |
| 800bp | Paired-end | 700 - 900 | 2 x 160 | 239,761,152 | 76,723,568,640 | 32.0 |
| 3-5kb | Mate-pair | 3K - 5K | 2 x 160 | 236,298,919 | 75,615,654,080 | 31.5 |
| 5-7kb | Mate-pair | 5K - 7 K | 2 x 160 | 258,583,949 | 82,746,863,680 | 34.5 |
| 10-15kb | Mate-pair | 10K - 15K | 2 x 160 | 268,922,922 | 86,055,335,040 | 35.9 |

*1.2. Rainbow trout de novo draft assembly*

We used the DeNovoMAGIC genome assembly pipeline from NRGene (nrgene.com;(Hirsch et al., 2016)) to generate a 2.17Gb genome assembly, containing 139,726 scaffolds with N50 greater than 1.67Mb. The assembly statistics are summarized in Supplementary Table 2.

***Supplementary Table S2.*** *Summary statistics of the genome scaffolds assembly.*

|  | Scaffolds | Contigs |
| --- | --- | --- |
| Number | 139,800 | 559,855 |
| N50 | 1,670,138 bp | 13,827 |
| Longest | 22,431,463 bp | 238,584 |
| Average | 13,787 bp |  |
| Shortest | 351 bp | 251 |
| Total Length | 2,178,999,613 bp |  |
| Total Ungapped Length | 1,927,483,719 bp |  |

*1.3. High-density linkage maps*

The construction of a dense linkage maps and its integration with a high-quality *de novo* draft assembly, provide the means to construct chromosome-length sequences for rainbow trout. A high-density linkage map for rainbow trout was constructed based on genotyping of the 57K SNP Axiom^®^ Trout Genotyping Array (Palti et al. 2015a) on in total 5,716 fish collected across 111 full-sib families from a commercial Norwegian population (AquaGen; N=3,108), 25 full-sib families from a commercial US company (Troutlodge, Inc.; N=1,949) and 10 full-sib families from the USDA-ARS-NCCCWA odd-year breeding population (N=659). Following quality control of raw genotype data as previously described (Palti et al. 2015a), linkage mapping was performed with Lep-MAP software (Rastas et al. 2013). First, SNPs were assigned to linkage groups with the ‘SeparateChromosomes’ command using increasing LOD thresholds until the observed number of linkage groups corresponded with the haploid chromosome number of 29. Additional SNPs were subsequently added to the groups with the ‘JoinSingles’ command at a more relaxed LOD threshold, and finally SNPs were ordered in each linkage group with the ‘OrderMarkers’ command. Numerous iterations were performed to optimise error and recombination parameters. A total of 46,174 SNPs were mapped to 29 linkage groups, with an average of 1,592 SNPs per group. The number of SNPs assigned to each group ranged from 739 to 2,851. The resulting male and female linkage maps span a total of 2,044 and 3,731 centimorgans (cM), respectively. Rainbow trout linkage groups were assigned to specific chromosomes following the nomeclature by Phillips et al., (2006).

In addition to the SNP-based linkage map, two Restriction-site Associated DNA (RAD) sequencing based linkage maps were used to anchor scaffolds to linkage groups. In the first RAD SNP map, 628 doubled haploid fish from a single cross of two doubled haploid lines (Whale-Rock x Swanson) were genotyped with 30,919 SNPs. The rQTL package was used to anchor and order the markers on 29 linkage groups following the methods of Miller et al. (2012). In the second RAD map, 701 fish from the 10 NCCCWA full-sib families were genotyped with 10,059 SNPs that were anchored to the 29 linkage groups by two-point linkage analysis with the Multi-Map software package following the methods as previously described (Palti et al., 2015b, Liu et al., 2015).

*1.4. Construction of chromosome sequences*

Sequence flanking each marker was used to precisely position all genetic markers to contigs and scaffolds in the rainbow trout *de novo* assembly (see Section 1.2) using megablast (Altschul et al. 1990), and thereby associate sequence with linkage groups. Altogether, we mapped 77,578 markers with linkage information to unique positions on 3,293 scaffolds containing a combined length of 1,781,543,814 bp (82% of the scaffolds assembly). The analysis revealed 36 chimeric scaffolds containing at least two markers from each of different linkage groups. If supported by comparative information from the Atlantic salmon genome (Lien at al. 2016), chimeric scaffolds were selectively ‘broken’ between incorrectly linked contigs. After breakage of scaffolds, linkage information was used, together with comparative information from the Atlantic salmon genome, to order and orient both broken and unbroken scaffolds within 29 chromosomes. Conserved synteny between rainbow trout and Atlantic salmon were determined by aligning Atlantic salmon chromosome sequences with rainbow trout scaffolds using LASTZ (Harris 2007). LASTZ command line script; --targetcapsule=LZ_target_capsule query.fa —nochain --gfextend --nogapped —identity=90.0..100.0 —matchcount=100 —format=general —rdotplot=plotoutput.txt.

Next, Dovetail Chicago library sequencing was used to validate, refine ordering and orientation of scaffolds and to integrate additional scaffolds within the existing chromosome sequences. DNA extraction, library preparation and sequencing and joining of scaffolds were done by Dovetail Genomics (<https://dovetailgenomics.com/>). A Chicago library was prepared as described previously (Putnam et al., 2016) using high quality genomic DNA extracted from 1 ml of blood collect from Swanson DH line fish at Washington State University following the university IACUC approved protocol. Briefly, 500ng of high molecular weight gDNA (~50kb mean fragment size) was reconstituted into chromatin *in vitro* and fixed with formaldehyde. Fixed chromatin was digested with DpnII, the 5’ overhangs were filled in with biotinylated nucleotides, and free blunt ends were ligated. After ligation, crosslinks were reversed and the DNA purified from protein. Purified DNA was treated to remove biotin that was not internal to ligated fragments. The DNA was sheared to ~350 bp mean fragment size, and a sequencing library was generated using NEBNext Ultra enzymes and Illumina-compatible adapters. Biotin-containing fragments were isolated using streptavidin beads before PCR enrichment of the library. The library was sequenced on an Illumina HiSeq 2500 in rapid run mode to produce 174 million 2 x 150bp read pairs for an estimated 62.8x physical coverage of the genome. Scaffolds from the rainbow trout assembly and Chicago library read pairs in FASTQ format were used as input data for HiRise, a software pipeline designed for using Chicago library sequence reads data to scaffold genomes (Putnam et al. 2016). Shotgun and Chicago library sequences were aligned to the rainbow trout assembly using a modified SNAP read mapper (<http://snap.cs.berkeley.edu>). The separations of Chicago read pairs mapped within draft scaffolds were analyzed by HiRise to produce a likelihood model for genomic distance between read pairs, and the model was used to identify putative wrong joins and score prospective joins. After scaffolding, shotgun sequences were used to close gaps between contigs. Notably, the Dovetail scaffolding assembly was only used to correct order and orientation of scaffolds only when supported by linkage data or by conserved synteny with the Atlantic salmon genome. Dovetail scaffolding separated 44 presumably chimeric scaffolds into smaller scaffold and joined 7,982 scaffolds into 3,991 larger scaffolds, including closing 99 gaps by adding or “filling-in” new sequence data. Moreover, it added 3,531 previously unmapped scaffolds (total of 38.5 million bases) to the chromosome sequences through joining or bridging to previously mapped scaffolds.

The final rainbow trout assembly (Omyk_1.0) contains 1.92 Gb of sequence assembled into 29 single chromosome sequences and 229 Mb of unplaced sequence. Omyk_1.0 is available at NCBI GenBank accession GCA_002163495.1.

**2. Post-Ss4R rediploidization characteristics**

To characterize the duplicated genome structure, and investigate mechanisms of rediploidization in different regions of the rainbow trout genome, we aligned rainbow trout chromosome sequences using LASTZ (Harris, 2007) to disentangle conserved collinear blocks of homeology.

The same approach was used to determine conserved synteny between with rainbow trout and Atlantic salmon. LASTZ command line script; --targetcapsule=LZ_target_capsule query.fa —nochain --gfextend --nogapped —identity=75.0..100.0 —matchcount=100 —format=general —rdotplot=plotoutput.txt. In total 88 blocks, amounting 1.74 Gb (90.6%) of chromosome sequences, were identified (Supplementary Table S3). The remaining sequence (181 Mb) could not be matched to a homeolog region with high collinearity suggesting that rearrangements, deletions, and/or fragmentation has preferentially eroded in some chromosome regions. The majority of these unmatched sequences are located at chromosome ends or at the sites where there is evidence for ancestral chromosome fusion (see red rectangles in Figure 1), suggesting that substantial amount of sequence has been lost in these regions.

Sequence similarity between homeologous sequences within the rainbow trout genome was determined in 1Mb intervals by averaging local percentage of nucleotide sequence identity using High-scoring Segment Pair (HSP) from LASTZ alignments (Harris, 2007). This revealed elevated sequence similarity (>90%) between blocks clustered within seven pairs of chromosome arms; 2p-3p, 6q-26, 7p-18p, 10q-19p, 12q-13q, 13p-17p and 15q-21p, corresponding perfectly with the regions displaying delayed rediploidization in the Atlantic salmon genome (Lien et al. 2016). Elevated sequence similarity, but not as extreme as for the seven highly homeologous regions, was found for 14q-25p and 24-27. These two regions have also elevated sequence similarity in Atlantic salmon (5p-9qb and 9qc-20qb). We note that an eighth region (1q-23) that has been found to have high sequence similarity in many *Oncorhynchus* species (Sutherland et al. 2016) does not show a similar pattern here. The rest of the chromosome arms in the rediploidized part of the rainbow trout genome contain larger rearrangements that prevent homeologous recombination (Lien et al. 2016). Together, this suggests a common rediploidization history empowered by larger rearrangements blocking homeologous recombination for the majority of the two genomes.

Another mechanism to facilitate rediploidization may be to obstruct homeologous recombination by flipping whole arms of metacentric chromosomes. Atlantic salmon and rainbow trout genomes possess an interesting system to investigate this because the two species have similar number of chromosomes (2N=59, range 54-64)   but represent very different karyotypes with eight metacentric chromosomes in Atlantic salmon (NF = 72-74) and 23 metacentrics in rainbow trout (NF = 104). To determine orientation of chromosome arms we aligned the chromosome sequences of the two species with the pike genome ([GCA_000721915.3](https://www.ncbi.nlm.nih.gov/assembly/954671)). This revealed nine chromosome arms in rainbow trout with likely flipped orientation to centromere for one of the nine chromosome pairs; 1p-2q, 4p-5q, 4q-8p, 5p-29, 7q-17q, 8q-28, 9p-21q, 10p-12p, 19q-25q in rainbow trout. Looking into the alignment of these chromosome arms (see Figure 1), five of the regions (1p-2q, 4p-5q, 4q-8p, 5p-29 and 10p-12p) contain large additional rearrangement that could prevent homeologous recombination. However, flipped orientation to centromere may be of more importance for preventing homeologous recombination for 7q-17q, 8q-28, 9p-21q, 19q-25q.

**3. High-density linkage mapping to resolve** **genome organization**

Detailed recombination rate estimates across the genome are instrumental for understanding the evolutionary history of genome organization and for investigating how changes in genome organization may lead to phenotypic and adaptive divergence (Stapley et al. 2017). The high-density linkage map (based on 44,910 markers and 5,716 fish) and chromosome-scale genome assembly presented herein allowed us to describe sex-specific recombination rates across the genome and characterize large structural variation in the rainbow trout genome.

First, we documented striking recombination differences between the sexes, with females recombining along whole chromosomes and male recombination strongly localized towards telomeric regions (Figure 1; Extended Data Figure S2a, c.; Allendorf and Danzmann 1997; Gharbi et al. 2006; Lien et al. 2011). Female recombination is clearly reduced at centromeres (Extended Data Figure S2a), which also correlates with centromere bias in repeat content (Figure 1). Male recombination is highly elevated at most telomeres, but at the same time is repressed at the telomeric regions of homeologous chromosome arms with high sequence similarity (Extended Data Figure S2b). We postulate that the latter is due to multivalent pairing at meiosis obstructing homolog recombination in these regions (Allendorf and Thorgaard 1984; Lien et al. 2011).

Second, karyotyping in rainbow trout has suggested a variable chromosome number (2N = 58-64)(Thorgaard, 1983), but not which chromosomes are involved in this variation. Here we use linkage mapping to identify these as Omy04, Omy14 and Omy25 (Figure 1; Supplementary Figure S1 d.-f.).

Third, linkage mapping revealed patterns of recombination interference in the linkage maps of Omy05 and Omy20, suggesting larger polymorphic rearrangements on these two chromosomes. In support of these findings we detected extended blocks of elevated linkage disequilibrium spanning the entire rearrangements, both in the pedigree and in wild populations. Haplotypes tagging of the various rearrangements were identified and utilized to classify the parents of the mapping families. Subsequent linkage mapping in parents being homozygous for distinct haplotypes disclosed the structure of the rearrangements, whereas linkage mapping in heterozygous parents documented almost complete repression of recombination across the rearrangements. For Omy05, we revealed two adjacent inversions of 22.83 and 32.94 Mb. The first inversion flips the centromere (pericentric), which may help to stabilize the rearrangement.

**4. Rainbow trout repeats**

*4.1. A repeat library for rainbow trout*

The rainbow trout repeat library creation process was guided by the methodology used for Atlantic salmon (Lien et al. 2016). In brief, repeats from both existing and *de novo* sources were processed in a software pipeline that incorporated repetitiveness validation, redundancy removal, non-transposable element (TE) host gene identification and classification steps. The final rainbow trout library contained 2,599 repeat sequences, of which 1,240 (47.7%) were classified.

*4.2. Source libraries and repetitiveness validation*

Source libraries consisted of previously reported repeat sequences from Atlantic salmon and rainbow trout (Berthelot et al. 2014; Lien et al. 2016), all Salmoninae repeats present in the RepBase database as of February 2016 (Bao et al. 2015), and a *de novo* repeat library created using the rainbow trout genome sequence contigs and RepeatModeler v1.0.8 (Smit and Hubley 2008). All of the sequences from source libraries were aligned to the rainbow trout genome using BLASTN (word_size 7; Altschul et al. 1990), and classified according to the number and length of the resulting HSPs. Any source library repeats generating at least three alignments which occupied over 80% of the query sequence with at least 80% sequence similarity were categorized as ‘High Confidence’ (HC). Sequences not designated as HC were classified as ‘Lower Confidence’ (LC) if they produced 10 or more alignments of at least 100 bp; otherwise they were eliminated. Many LC sequences failed to generate longer HC-quality alignments due to an internal stretch of ambiguous (N/X) bases, or because they were the result of an inappropriate concatenation of different repeat elements. To ameliorate the presence of such confounding sequences, all LC sequences were split wherever the number of long (80+ bp) BLASTN HSPs overlapping a particular query sequence base dropped below 10 over 10 consecutive bases; low coverage sequence was removed.

*4.3. Redundancy removal*

Validated sequences from the source libraries were merged and redundant sequences were removed in accordance with the 80-80-80 rule proposed by Wicker et al. (2007). First, HC sequences were aligned against each other using BLASTN in an all-by-all manner. A sequence was identified as redundant and removed if there existed HSPs between it and another, longer sequence which: i) were at least 80 bp long; ii) had a similarity of at least 80%, and; iii) together occupied at least 80% of the shorter sequence, with no HSPs overlapping by more than 15 bp. Following this HC library-only procedure, analogous processes were performed to reduce redundancy in the LC library on its own, and then to merge the LC library into the HC library. At this point, the set of 48 manually-curated Tc1-Mariner TE sequences described below were also merged into the combined library.

*4.4. Non-TE host gene detection*

In order to identify non-TE host genes that should be removed from the final library, all repeat sequences were compared to two protein databases using BLASTX: REPET-formatted RepBase v20.05 and the January 26^th^, 2017 version of SwissProt UniProtKB (Apweiler et al. 2004). Repeat library sequences were categorized as non-TE host genes and removed if they possessed a higher-scoring alignment to a non-TE sequence in the UniProtKB database (with E-value ≤ 1e^-10^) than they did to a sequence in the REPET-formatted RepBase database. During the ensuing repeat classification step, rDNA genes were also identified (and removed) with the PASTEClassifier.py tool (Flutre et al. 2011; Hoede et al. 2014), which made use of eukaryotic rRNA sequences from release 115 of the SILVA rRNA database (Yilmaz et al. 2013).

*4.5. Repeat classification*

Repeat sequences were classified in accordance with the taxonomy of Wicker et al (2007) based on both their similarity to reference TEs and structural information. First, sequences were classified to the superfamily level if they generated a ‘good’ BLASTN alignment to a sequence in the REPET-formatted RepBase TE nucleotide library (v20.05). Good alignments were considered to be those that: i) were at least 80 bp long; ii) possessed at least 80% sequence similarity between query and subject, and iii) occupied at least 80% of the query repeat. If no BLASTN-based classification was found for a sequence, it was aligned to the REPET-formatted RepBase TE protein library (v20.05) using BLASTX, where superfamily was assigned based on the best HSP with E-value less than 1e^-10^. The PASTEClassifier.py tool was used to facilitate further classification, and made use of reference databases including the aforementioned SILVA rRNA database and a REPET-formatted set of PFAM HMM Gypsy profiles (v26.0) available on the REPET website (<https://urgi.versailles.inra.fr/Tools/REPET)>. Otherwise unclassified sequences flagged as miniature inverted-repeat TEs (‘MITEs’) by PASTEClassifier were assigned to the TIR order, while any sequences flagged as ‘PotentialChimeric’ were manually reviewed and, if an obvious category couldn’t be identified, were labelled ‘Unknown’. Finally, all sequences were reviewed by dotplot using the Geneious software (Kearse et al. 2012), and those which consisted predominantly of a tandem repeat (satellite) or simple repeat sequence were classified as such.

*4.6. Repeat abundance in rainbow trout*

The abundance of different repeat types in the rainbow trout genome was assessed using RepeatMasker; the results are detailed in Supplementary Table S5. The results are largely consistent with those found for Atlantic salmon (in which 59.89% of the genome was determined to be repeat-derived), and the large number of superfamilies is consistent with what has previously been noted for actinopterygian fishes (Chalopin et al. 2015).

*4.7. Creating a library of curated Tc1-Mariner sequences*

A library of manually-curated Tc1-Mariner consensus sequences was created in order to facilitate a more in-depth analysis of what is by far the most abundant superfamily in the rainbow trout genome. Putative Tc1-Mariner sequences from a preliminary repeat library were aligned against the rainbow trout genome using BLASTN, and up to 200 random copies were extracted for each repeat, along with up to 2,000 bp of flanking sequence. These sequences were aligned using MUSCLE v3.8.31 (Edgar 2004) and the boundaries of full-length Tc1-Mariner elements were visually identified and trimmed. Following manual review of the resulting trimmed alignments, consensus sequences were generated, aligned using MUSCLE, and combined into a single Neighbour-Joining tree (Jukes-Cantor method). This tree was used to identify and remove redundant consensuses. In total, 48 Tc1-Mariner consensus sequences were identified in rainbow trout. When merged with the 40 sequences that were previously identified in Atlantic salmon (Lien et al. 2016), a total of 60 salmonid Tc1-Mariner sequences were found. NJ trees for all families evinced the starburst-like topology which is characteristic of Tc1-Mariner activity in many genomes (Feschotte and Pritham 2007; Pace et al. 2008).

*4.8. Visualizing historical Tc1-Mariner activity in salmonids*

In order to determine the relative origin of different Tc1-Mariner elements that are present in the modern rainbow trout genome, the percent similarity between aligned members of individual Tc1-Mariner families was used as a proxy for time. In this paradigm, pairs of sequences which were created longer ago would exhibit a larger amount of sequence divergence between them than would copies that arose more recently. To create a timeline of abundance and activity for three salmonid species (Extended Data Figure S1a) and one solely for rainbow trout (Extended Data Figure S1b), up to 200 randomly-selected TE copies were obtained for each Tc1-Mariner family, aligned with MUSCLE, and the percent sequence similarity between each pair of aligned sequences was calculated. All TE copies were required to be at least 60% of the length of their family consensus to facilitate alignment, and were confirmed to belong to the family to which they were attributed using a reciprocal-best BLASTN hit (RBH) strategy.

Combined Tc1-Mariner abundance and activity period estimates were created using the 60 families identified in Atlantic salmon and rainbow trout for the genomes of Atlantic salmon (GCF_000233375.1), rainbow trout (GCF_002163495.1) and Chinook salmon (GCA_002872995.1); these estimates are visualized in Supplementary Figure S1. Many families show differential activity between the three salmonid species, however one family (omyk_TCE_37) is notably prolific in rainbow trout. This family has been active recently in salmonid evolutionary history and makes up more than 2% of the entire rainbow trout genome; it is much less abundant in the other two species. An activity timeline of 48 Tc1-Mariner families identified in rainbow trout (Extended Data Figure S1b) was created by scaling density plots of the percent similarity values for each family by their abundance in the genome, and as with Supplementary Figure S1 also reveals a substantial recent burst of Tc1-Mariner activity.

**Table S5. Repeat abundance in the rainbow trout genome.** Percentage values are based on the number of bases in assembly GCF_002163495.1, excluding tracts of ≥ 20 ambiguous (N/X) bases (1.93 Gbp total). Values for individual TE categories are summed from annotations extracted from the RepeatMasker *.out file, which are not necessarily additive (ie, repeats can occasionally overlap each other).

**Table S5**

| **Repeat Type** | **Order** | **Superfamily** | **% Genome** |
| --- | --- | --- | --- |
| **Class I TEs** | **All** | **All** | **17.83** |
|  | **LTR** | **All** | **2.76** |
|  |  | Gypsy | 2.13 |
|  |  | ERV | 0.42 |
|  |  | Copia | 0.11 |
|  |  | Bel-Pao | 0.10 |
|  | **DIRS** | **DIRS** | **0.11** |
|  | **PLE** | **Penelope** | **0.17** |
|  | **LINE** | **All** | **14.51** |
|  |  | CR1 | 0.03 |
|  |  | Crack | 4.49 |
|  |  | Hero | 0.07 |
|  |  | L1 | 0.65 |
|  |  | L2 | 1.74 |
|  |  | Nimb | 0.21 |
|  |  | R2 | <0.01 |
|  |  | Rex1 | 5.40 |
|  |  | RTE | 0.01 |
|  |  | RTEX | 1.13 |
|  |  | Tx1 | 0.77 |
|  | **SINE** | **All** | **0.28** |
|  |  | tRNA | 0.17 |
|  |  | Deu | 0.10 |
| **Class II TEs** | **All** | **All** | **23.49** |
|  | **TIR** | **All** | **20.40** |
|  |  | CMC-EnSpm | 0.21 |
|  |  | Ginger | 0.05 |
|  |  | Harbinger | 0.10 |
|  |  | hAT | 2.45 |
|  |  | IS3EU | 0.07 |
|  |  | ISL2EU | 0.01 |
|  |  | Kolobok | 0.04 |
|  |  | PiggyBac | 0.40 |
|  |  | Sola | 0.04 |
|  |  | Tc1-Mariner | 17.04 |
|  | **Dada** | **Dada** | **0.00** |
|  | **Crypton** | **Crypton** | **0.17** |
|  | **Maverick** | **Maverick** | **0.06** |
| **Unclassified** |  |  | **14.29** |
| **Satellites** |  |  | **0.98** |
| **Simple repeats** |  |  | **1.93** |
| **Low-complexity** |  |  | **0.36** |
| **Total Masked** |  |  | **57.13** |

**5. Positioning of sdY in the rainbow trout genome**

A previous study has positioned the rainbow trout sex-determining locus, *sdY,* within a 31 kb fragment (KJ851798.1) containing highly transposon-like sequences (Faber-Hammond et al., 2015). This sequence aligns well to a 65.5 kb unmapped scaffold in the *de novo* rainbow trout draft assembly (scaffold241259_0-65500_unkn). To tie this scaffold into chromosome sequence we used information from Phillips et al. 2013, who characterized a 800 kb BAC contig containing sdY, the male-specific genetic marker OmyY1, as well as a number of other protein coding genes. Three of these genes, KC686348.1, KC686346.1 and KC686351.1, map to a 452 kb scaffold (scaffold27334_0-452001_omy29) located at approx. 5 Mb on Omy29, which substantiates adjacent location of the two scaffolds.

**6. Omy05 re-arrangements and diversity analysis**

**6.1. Whole genome re-sequencing**

Genomic DNA was extracted from fin clips of 61 rainbow trout. Whole-genome paired-end sequencing libraries were prepared and sequenced using the Illumina HiSeq 2000 or 2500 platforms providing an average of 15X and minimum of 8X genome coverage per sample. The sequenced samples included 11 clonal lines from Washington State University, 38 steelhead and resident rainbow trout from wild and hatchery-origin populations distributed throughout the native range of the species, and 12 fish from the AquaGen rainbow trout aquaculture breeding program. The population of origin and geographic location of each sample are given in Supplemental Table S4. A total of 31,441,105 SNPs was identified genome-wide, of which a subset of SNPs that were mapped to chromosome Omy05 was further used for population genetic and functional analyses as described below.

**6.2 Population Genetic Analysis of Resequencing Data**

A subset of 31 individuals of known geographic origin that were known to be homozygous for either “A” or “R” MAR genotypes (Figure 2c ; Table S4) was extracted from the SNPeff data file and separated into genome-wide and omy05 inversion data sets with *vcftools* v. 0.1.14. Sequence diversity (π) and Weir and Cockerham’s *F_ST_* were calculated using *vcftools* and plotted in R using base R and *ggplot2*.A random subset of 40,000 SNPs was obtained with the shuf GNU coreutil for each data set. SNPs were analyzed in R v. 3.3.1 with Rstudio v. 1.0.136 using vcfR v. 1.4.0 (Knaus et al. 2017) and *adegenet* v. 2.0.1 (Jombart et al. 2008; Jombart and Ahmed 2011) packages visualized with *ggplot2* v. 2.2.1. Files in vcf format were imported into R with the read.vcfR function of vcfR and converted to *genind* objects for analysis with *adegenet* with the vcfR2genind function of vcfR. A PCA was conducted for each set of 40,000 SNPs from outside the inversion region and within it with the *dudi.pca* function of *ade4* v. 1.7-5.

**7. Fish movement experiment in Big Creek**

Across all samples, the frequency of *AA* genotypes was dramatically reduced among larger individuals of both sexes, while *RR* genotype individuals increased proportionally (Figure 3C). Sex ratio also shifted, from 50:50 among the youngest individuals to a strong male bias among larger residents (81.3% males >180 mm; Figure 3B), consistent with theoretical expectations and prior observations (Rundio et al. 2012; Ohms et al. 2013). Conversely, a significant female bias was detected among individuals identified as migrants (56%, p < 0.05). These results are independent of family or population structure, and the observed distribution of family sizes supports a positive interaction between the effects of Omy05 genotype and parental fitness (data not shown).

*7.1. Emigration Analysis*

**Table S6. Model selection for probability of detection by PIT tag antenna near stream mouth.**

| Model | No. smooths | AIC | ΔAIC |
| --- | --- | --- | --- |
| emigration ~ f(size at release, sex, genotype with complete sex-dependent dominance) | 4 | 1130.8 | 0.00 |
| emigration ~ f(size at release, sex, genotype with anadromous dominance) | 4 | 1135.7 | 4.90 |
| emigration ~ f(size at release, sex) | 2 | 1137.4 | 6.59 |
| emigration ~ f(size at release, sex, genotype with indepentent dominance) | 6 | 1146.1 | 8.04 |
| emigration ~ f(size at release, sex, genotype with resident dominance) | 4 | 1141.4 | 10.54 |
| emigration ~ f(size at release, genotype) | 3 | 1144.1 | 13.30 |
| emigration ~ f(size at release) | 1 | 1148.7 | 17.92 |

*7.2. Population, Family, and Fecundity Effects*

The observed association between Omy05 genotype and migratory tendency could be confounded by the presence of subpopulations of partially reproductively isolated resident and anadromous *O. mykiss* (Narum et al. 2004; Pearse et al. 2009). To investigate this, we evaluated the 92 SNP loci not linked to the Omy05 using the program STRUCTURE (Pritchard et al. 2000) to detect genetically distinct groups. We found no evidence for genetically distinct subpopulations of individuals within the sampled area based on these presumably neutral genetic markers (Figure ED10).

The results could also be confounded by the presence of a few large families if few parents produced many of the fish sampled in the study. To address this, we used the same dataset with the program COLONY(Jones and Wang 2010; Wang 2004) to identify full-sibling families among the 1,439 juveniles sampled in Fall 2009, which represent the majority of the study individuals and those most likely to have siblings in the sample. We conservatively considered only full-sibling families with at least five sampled members, since COLONY may incorrectly infer full-sibling groups with fewer individuals. We identified 103 full-sibling families with N ≥ 5 sampled individuals (range 5-39; Figure ED11), and these sibships were produced by 46 + 49 = 95 unique parents. While we cannot determine individual parental sexes, this large number of families and contributing parents suggests that the model results are not biased.

The results from the capture-recapture analyses show that AA and/or AR females are more likely to be anadromous than *RR* females, with associated differences in expected fecundity. This leads to the predictions that 1) the frequency of families, and 2) the distribution of family sizes, should reflect the maternal life-histories predicted by the Omy5 genotypes present in the offspring of a given family. To evaluate this, we compared the sizes of families with different Omy05 genotypes and their frequency of occurrence among the 30 full-sibling families for which all sampled offspring had a single Omy05 genotype (i.e. RR, AR, or AA offspring resulting from RRxRR, RRxAA, and AAxAA matings, respectively). Exclusively RR families were extremely rare and had few sampled individuals, consistent with the scarcity of adult female residents captured (N=11 females >180mm FL) and their low expected fecundity (Hendry et al. 2004), while exclusively AR and AA families were both larger and more common, as predicted by their likely anadromous maternal origins (mean(number of families): 6(2), 9.1(15), and 8.4(13), for RR, AR, and AA families, respectively, NS). Similarly large and abundant families were observed among the 59 full-sibling families with two MAR genotypes represented among the offspring (i.e. RR and AR, 10.7(20) or AR and AA, 9.0(39), resulting from RRxAR and AAxAR matings, respectively), as expected if the majority of both types of families come primarily from large, AR or AA, anadromous mothers. These patterns provide a direct link between adult phenotype and offspring genotype frequencies and highlight the dynamic interaction between the direct genetic effects of the Omy5 rearrangement and indirect maternal effects that contribute to variation among offspring in expression of anadromy driven by egg size, lipid content, or other differences between resident and anadromous mothers.
