## Supplementary material for "Sex-dependent dominance maintains migration supergene in rainbow trout": Pearse_etal_Additional_Figures

**Extended Data Figure S1a**

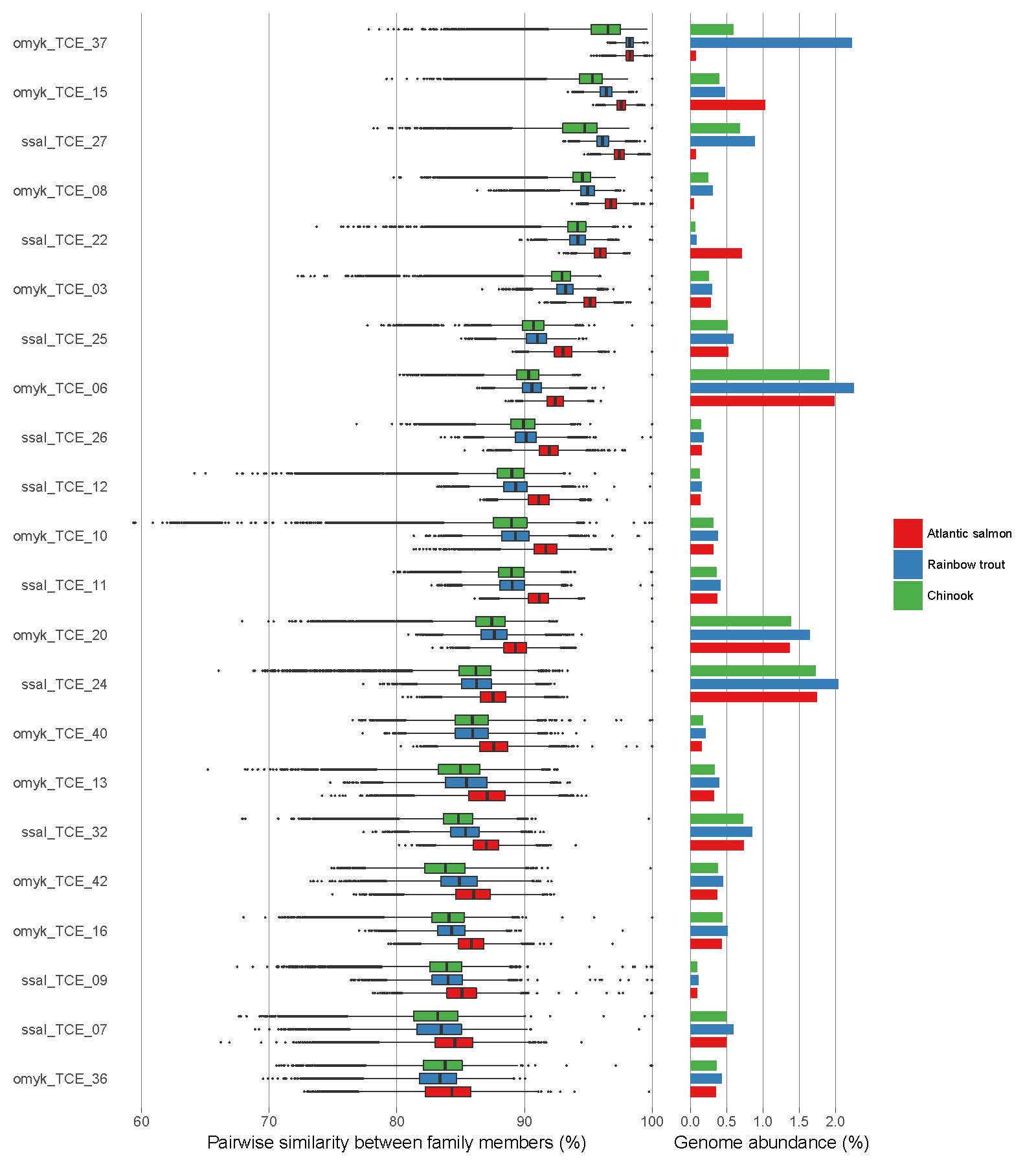

**Extended Data Figure S1a. Activity periods and abundance of Tc1-Mariner families in Atlantic salmon, rainbow trout, and Chinook salmon.** Lower sequence similarity between family members indicates a more ancient family. Activity and abundance are generally consistent among the three species until the time corresponding to ~93% sequence similarity, after which substantial differences in activity have occurred in concert with salmonid lineage divergence. Tc1-Mariner families displayed were identified in Atlantic salmon and rainbow trout and occupied at least 0.1% of the genome in one of the three species.

**Extended Data Figure S1b**

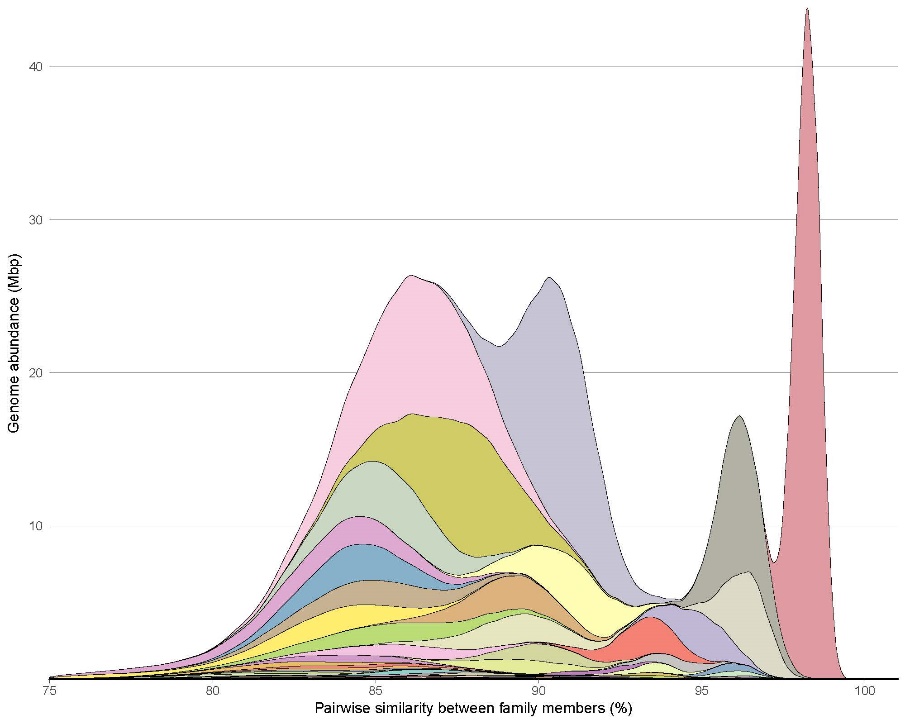

**Extended Data Figure S1b. Historical proliferation of Tc1-Mariner elements in the rainbow trout lineage.** Stacked density plot of pairwise similarity between Tc1-Mariner family members. The large initial peak with a maxima at ~86% corresponds roughly to the same time that the salmonid-specific whole-genome duplication took place. In the time corresponding to more than 93% similarity, differences in activity begin to appear between Atlantic salmon and rainbow trout in accordance with their ancestral divergence (compare to Figure 3a in Lien et al, 2016).

**Extended Data Figure S2**

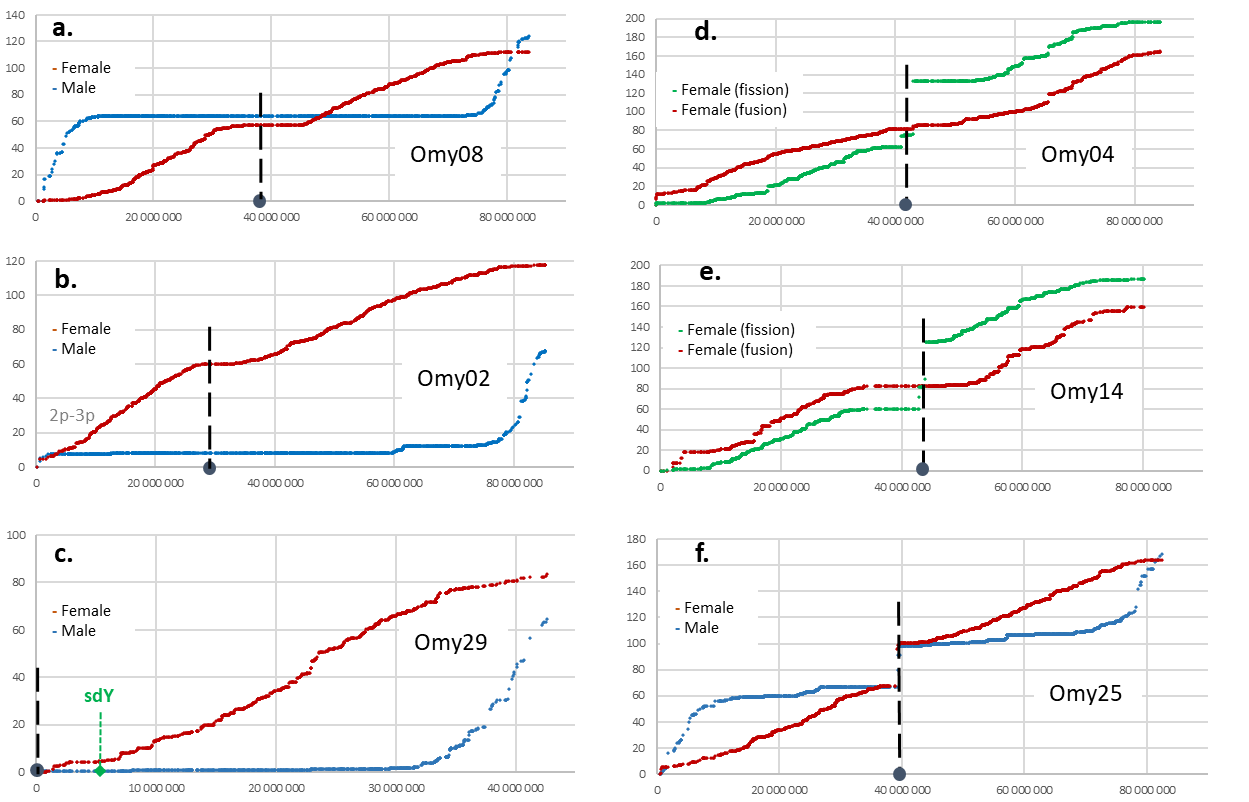

**Supplementary Figure S2. High-density linkage maps describing characteristic sex-specific recombination patterns and resolving variable chromosome numbers associated with centric fusions or fissions in rainbow trout. a.** Linkage map for the metacentric rainbow trout chromosome 8 (Omy08) demonstrating male recombination strongly localized towards both telomeres and female recombination repressed at the centromere. **b.** Linkage map for the metacentric rainbow trout chromosome 2 (Omy02) demonstrating elevated male recombination towards the telomeric region at the q-arm but repressed recombination at the p-arm typified by showing high sequence similarity to Omy03p. **c.** Linkage map for the acrocentric rainbow trout chromosome 29 (Omy29) with the sex-determining gene sdY located around 5 Mb demonstrating repressed male recombination for most of the chromosome except the telomeric region. **d.-f.** Rainbow trout chromosomes with variable chromosome numbers associated with centric fusions or fissions. Gaps in the linkage map at the centromere of Omy04, Omy14 and Omy25 are caused by fissions splitting metacentric chromosomes into two acrocentric chromosomes in some families.

**Extended Data Figure S3**

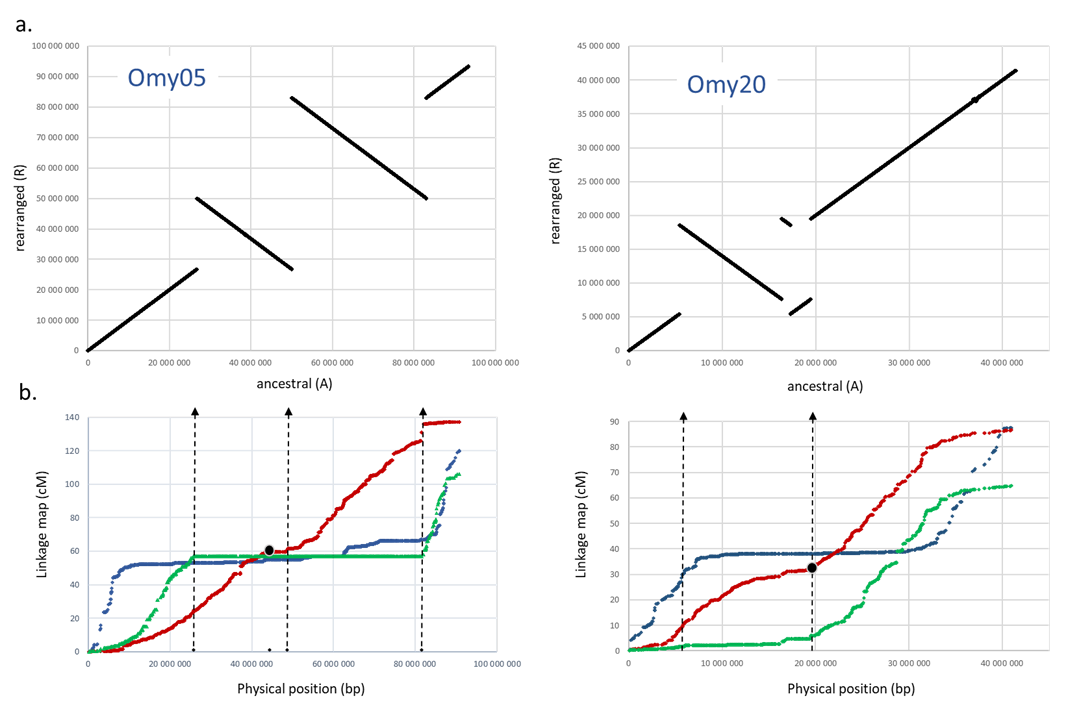

**Extended Data Figure S3. Large polymorphic chromosomal rearrangements in the rainbow trout genome.** Ancestral (A) and rearranged (R) chromosomal structures on Omy05 and Omy20. **a.** Ancestral (A) and rearranged (R) chromosomal structures on Omy05 and Omy20. **b.** Genetic linkage map constructed for parents with alternate Omy05 and Omy20 haplotypes. Red line; female map for homozygous ancestral (AA) parents. Green line; female map in heterozygous parents (AR). Blue line; male map for homozygous ancestral (AA) parents.

**Extended Data Figure S4**

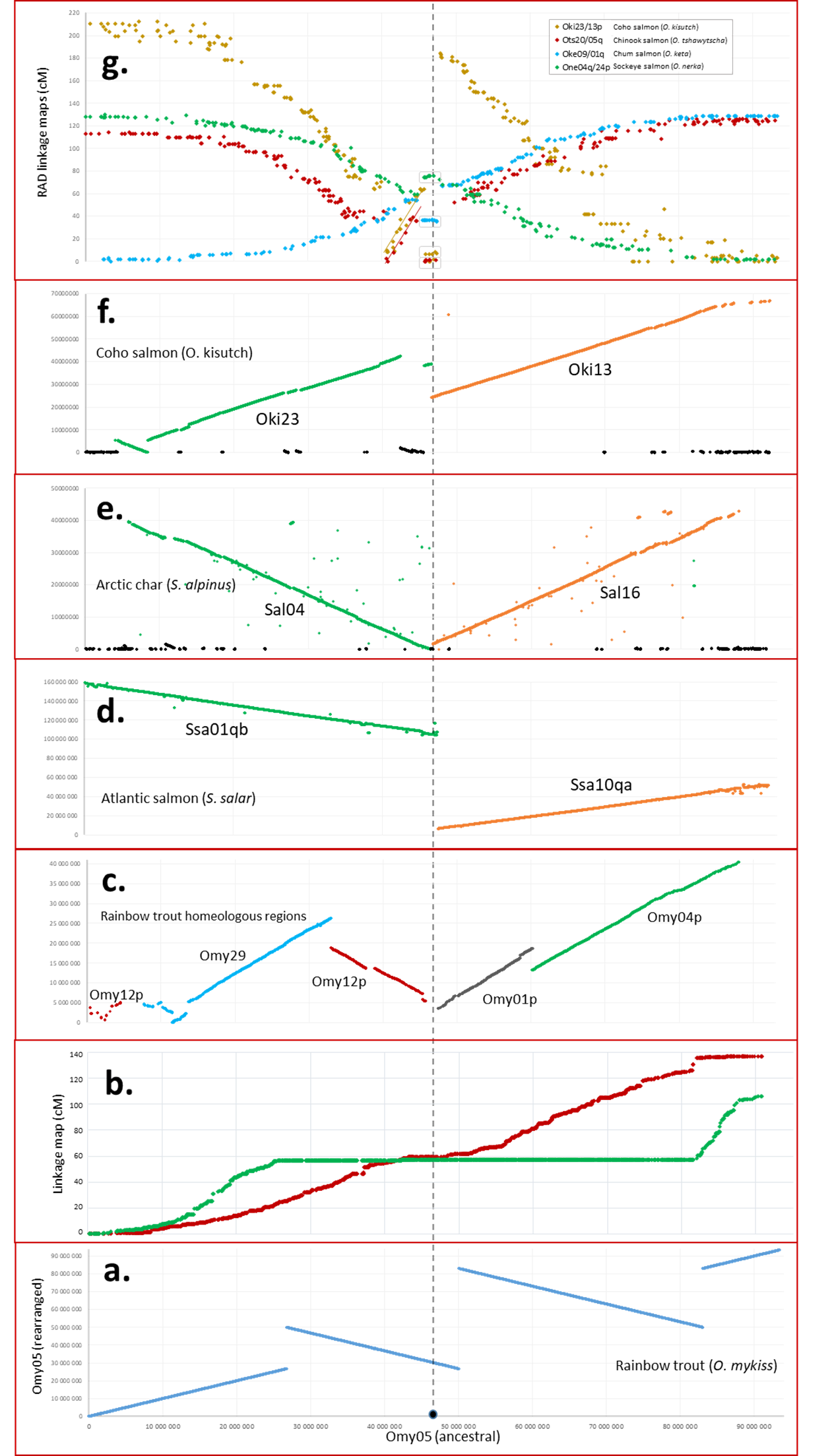

**Extended Data Figure S4. Chromosomal rearrangements on rainbow trout chromosome 5 (Omy05) and conserved synteny with other salmonid species. a.** Ancestral (A) or rearranged (R) haplotypes on Omy05 characterized by two adjacent inversions of 22.83 and 32.94 Mb. **b.**The alignment of Omy05 with the rest of the rainbow trout genome assembly show conserved collinear blocks of homeology with Omy12p, Omy29, Omy01p and Omy04p. **c.** The alignment of Omy05 with Atlantic salmon genome assembly (GCF_000233375.1) identifies highly conserved synteny with salmon chromosomes 1 and 10 (Ssa01qb and Ssa10qa). **d.** Alignment with the Arctic char genome (GCF_002910315.2) detects highly conserved synteny with char chromosomes 4 and 16 (Sal04 and Sal16). **e.** Alignment of Omy05 with coho salmon chromosome sequences (GCF_002021735.1) reveals conserved synteny with chromosomes 23 and 13 (Oki23 and Oki13). **f.** Comparison of Omy05 with RAD-based linkage maps for other pacific salmon reveal a smaller fragment at centromere of Omy05 which is rearranged in coho, Chinook, chum and sockeye compared to rainbow trout, Arctic char and Atlantic salmon, and a larger rearrangement that differentiate coho and chinook from the other salmonid species.

**Extended Data Figure S5**.

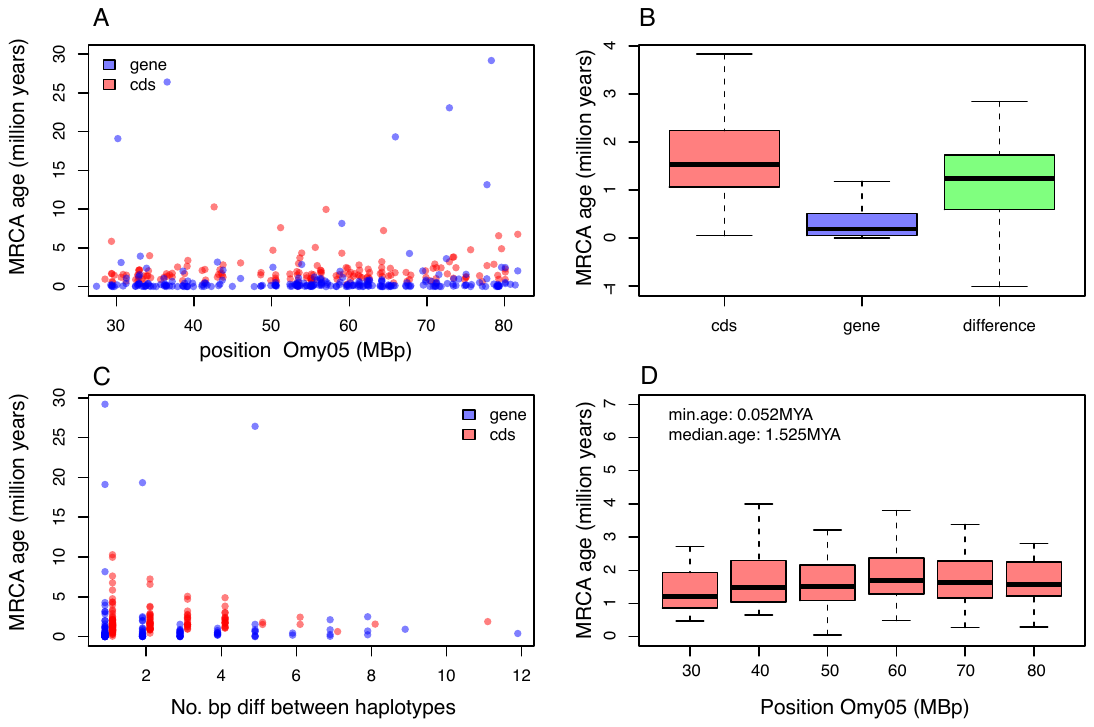

**Extended Data Figure S5. Age estimates of the Omy05 inversion complex.** A) Age estimates from individual gene and CDS across the inversion showing lack of a strong pattern associated with inversion break points. B) Boxplot of age estimates for all CDS, genes that passed filtering, and the difference between the two estimates. Older estimates and wider confidence intervals are obtained from CDS than those based on gene sequences. C) Plot of the number of base differences between haplotypes for gene and CDS alignments and its effect on age estimation. Despite potentially having more variants because of the inclusion of introns, the gene alignments actually have fewer base differences per haplotype on average owing to the removal of poor quality intronic alignments by Gblocks. As a result, the CDS based alignments provide more informative alignments for dating the inversion complex. D) Estimate of inversion age in 10Mb windows across the inversion complex for CDS estimates.

**Extended Data Figure S6**

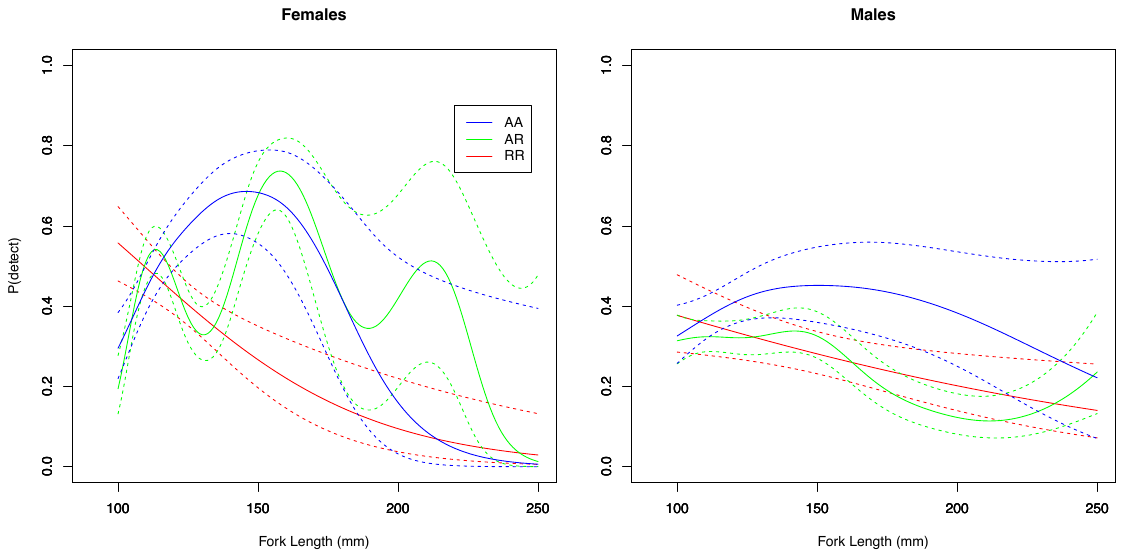

**Extended Data Figure S6**. Full model results from antennae detection emigration model in Big Creek, showing differential size-dependent migration of males and females with AR and AA, and RR genotypes, peaking at ~150mm.

**Extended Data Figure S7**

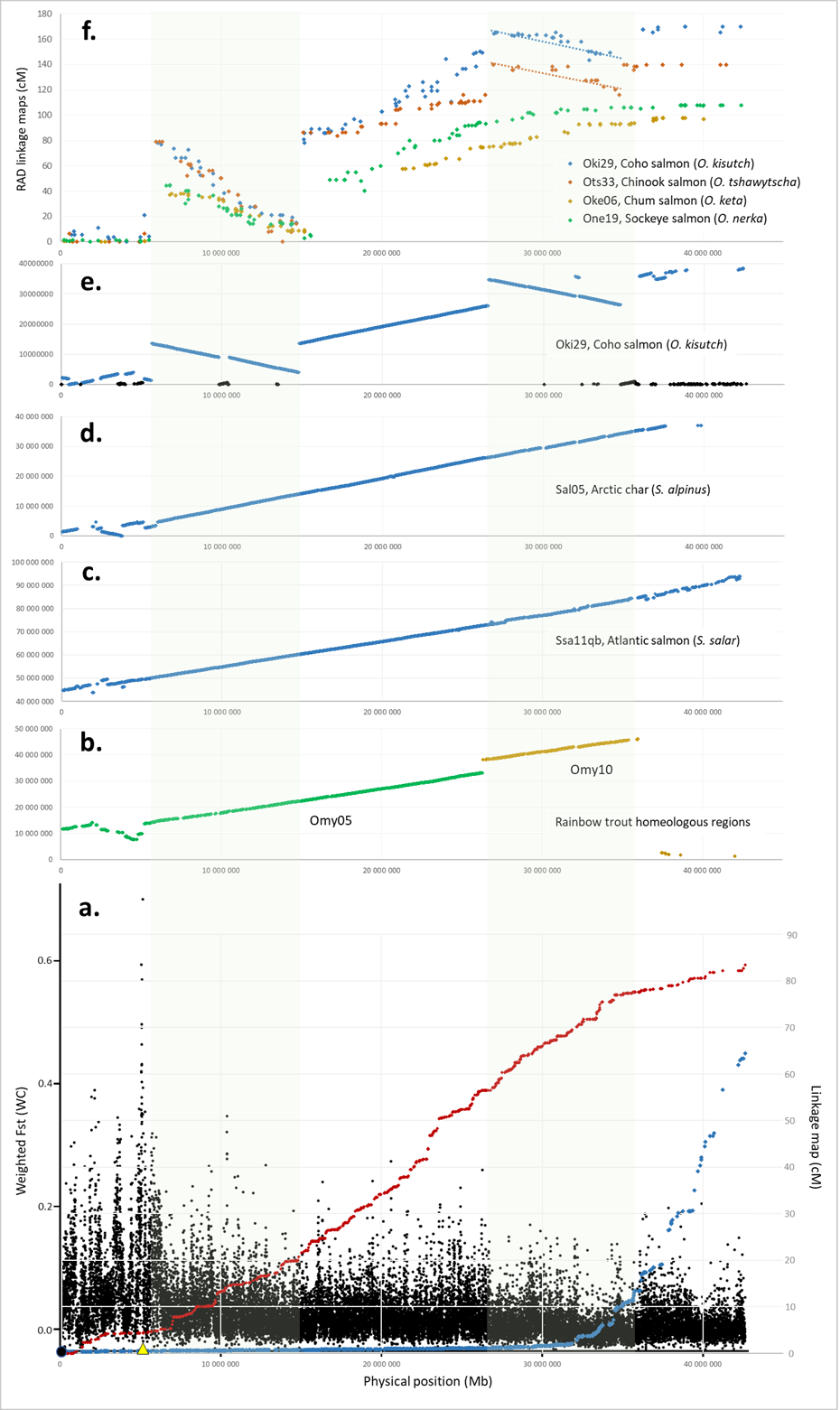

**Extended Data Figure S7. Linkage maps and conserved synteny with other salmonids for rainbow trout chromosome 29 (Omy29). a.** Sex-specific linkage maps for Omy29 (red dots; female, blue dots; male) show repressed recombination in males for the majority of Omy29. Genetic differentiation between males and females (Fst) is peaking at the *sdY* locus located at 5 Mb (yellow triangle) but is also elevated in the region between *sdY* and the centromere (Black Dot at 0 Mb). **b.** Regions of the rainbow trout genome homeologous with Omy29. Comparative genome sequence maps show highly conserved synteny with **c.** Atlantic salmon chromosome 11qb (ssa11qb) and **d.** Arctic char chromosome Sal05. **e.** Comparative mapping with coho salmon genome sequence reveals two larger rearrangements, one at 10 Mb and one at 30 Mb. **f.** Comparison of Omy29 with RAD-based linkage maps for other pacific salmon show that the rearrangement at 10 Mb is conserved among coho, chinook, chum and sockeye and differentiated from rainbow trout and Atlantic salmon.

**Extended Data Figure S8**

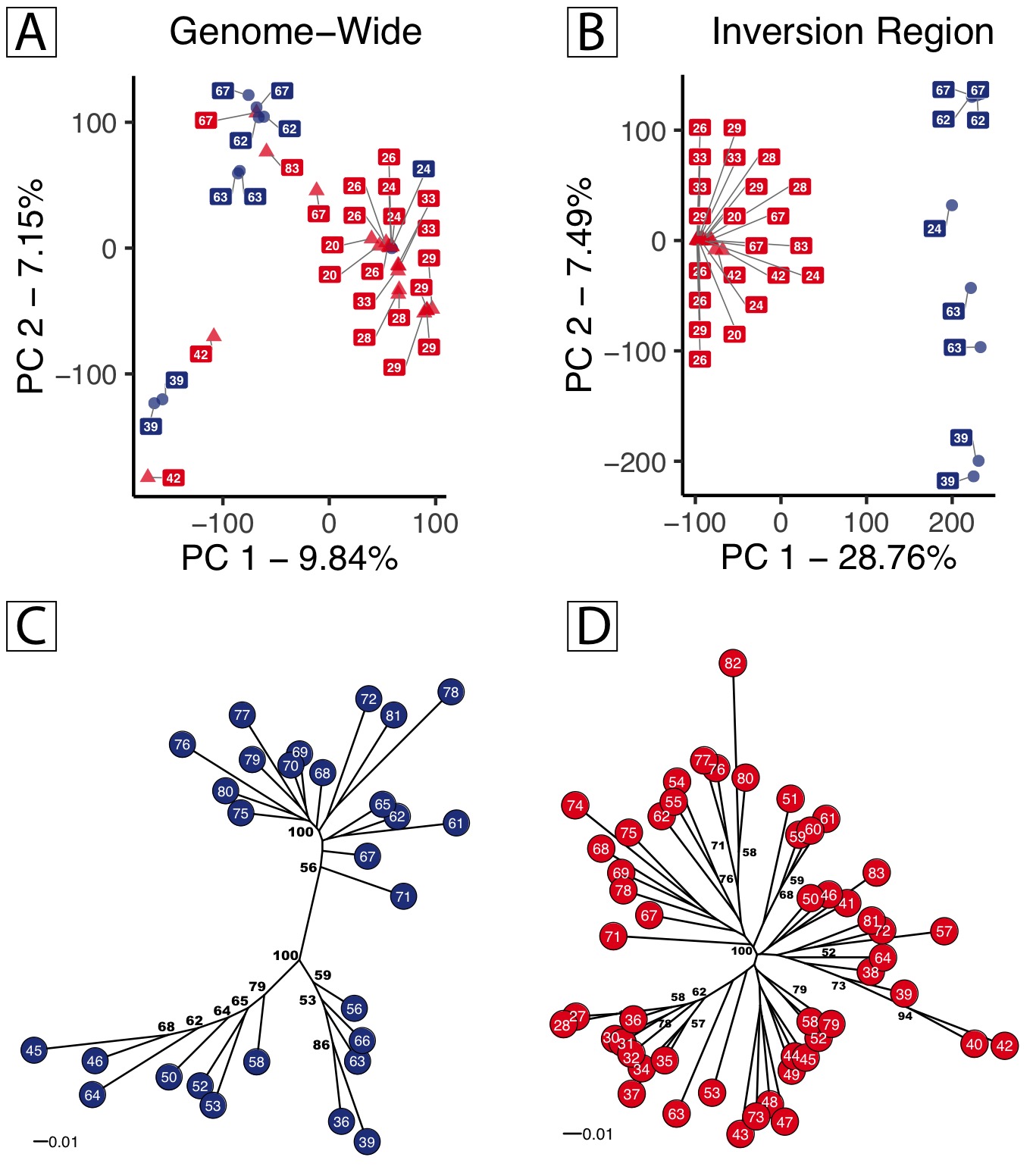

**Extended Data Figure S8: Genetic Diversity and Interrelationships of AA and RR Genotypes. –** Principal component (PC) analysis of genetic diversity estimated from whole-genome sequence data genome-wide (**A**) and within the chromosome Omy5 rearrangement (**B**). For both, a random subset of 40,000 SNPs were examined from homozygous AA and RR individuals of known geographic origin and plotted with the principal components containing the most variance. Homozygous AA individuals are plotted as blue circles with homozygous RR individuals represented by red triangles, with numbers indicating geographic locations corresponding to Figure 2d and Supplemental Information Table S4. While geographic structuring is apparent in the genome-wide SNP dataset, the inversion region separates clearly between RR and AA types with PC1 (28.76% of variation), and the subsequent second PC (7.49% of variation) corresponding to diversity within AA types. Similarly, population-level Neighbor-Joining trees of AA individuals and RR individuals from SNP survey data of homozygous AA individuals (**C**) and homozygous RR individuals (**D**) from sampled populations are depicted in a population level tree generated through chord distances. Support values for nodes with >50% bootstrap support generated from 1,000 bootstrap replicates are indicated. Population numbers correspond to Figure 2d and Supplemental Information Table S4.

**Extended Data Figure S9 – Inversion Frequency as a Function of Mean Monthly Temperature.** For each month of the year below barrier North American populations of rainbow trout are plotted as mean monthly temperature (*x* – axis) and inversion frequency (*y –* axis). Points are sized proportionally to sample size (N). A weighted least squares regression is depicted with the adjusted R^2^ value for each month of the year.

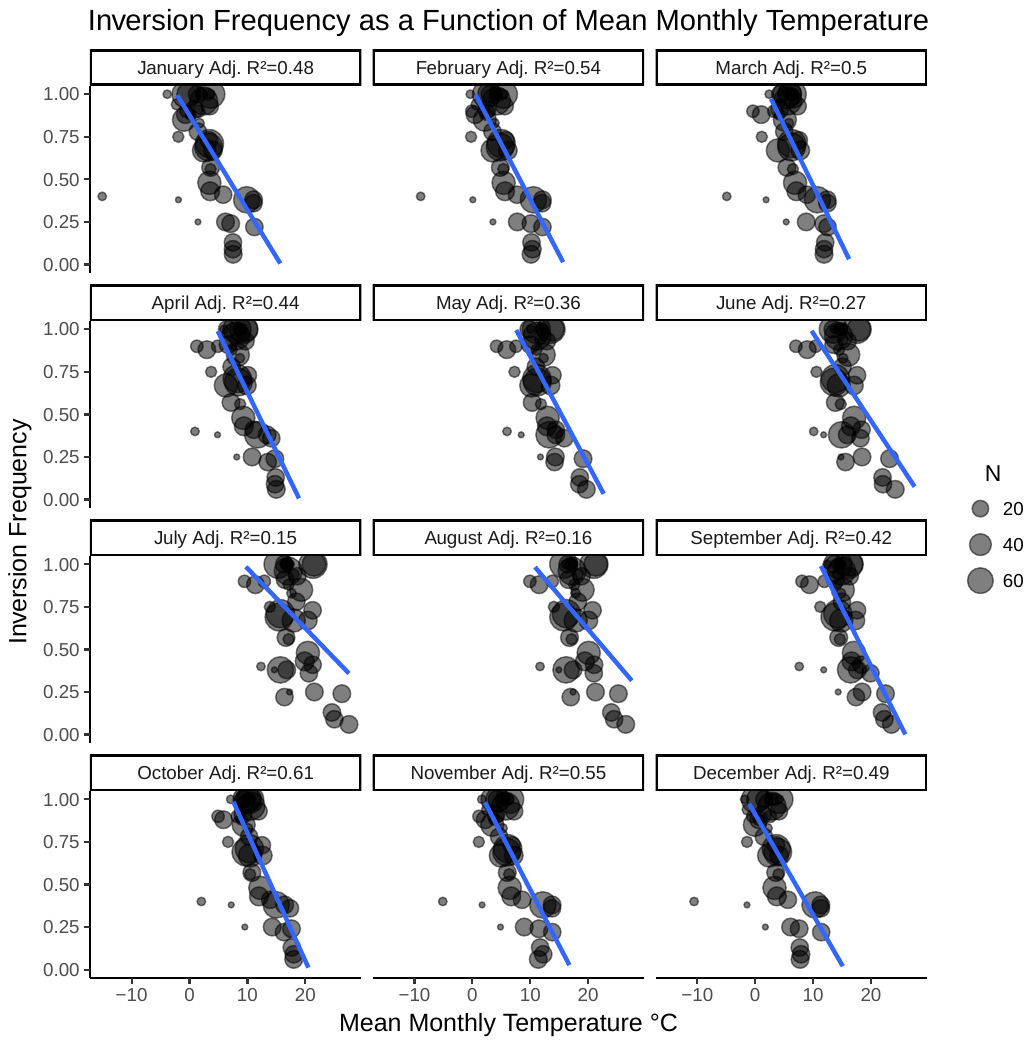

**Extended Data Figure S10** A) Graphical hypothesis of *Oncorhynchus mykiss* life-cycle showing differences in development, fecundity, and relative expression of alternative life-history patterns based on environment (temperature), sex, and Omy05 karyotype, leading to the alternative individual migratory life-histories *rainbow trout* and *steelhead*. B) Anadromous female steelhead and resident male rainbow trout. Photo: Morgan Bond.

A)

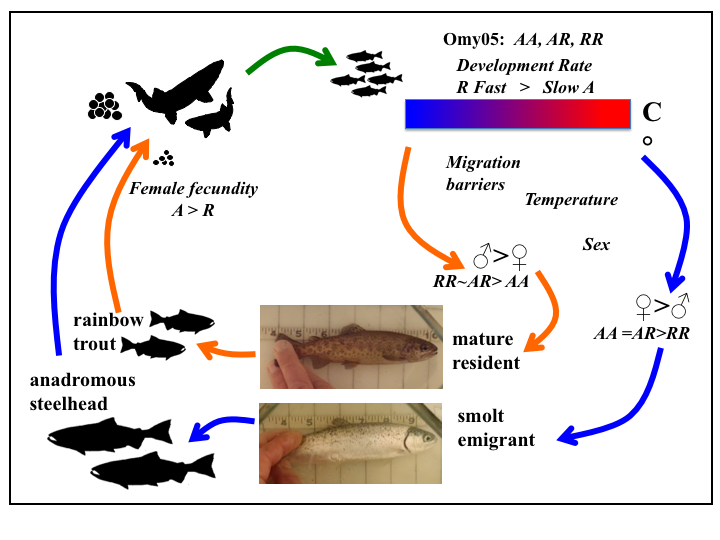

B)

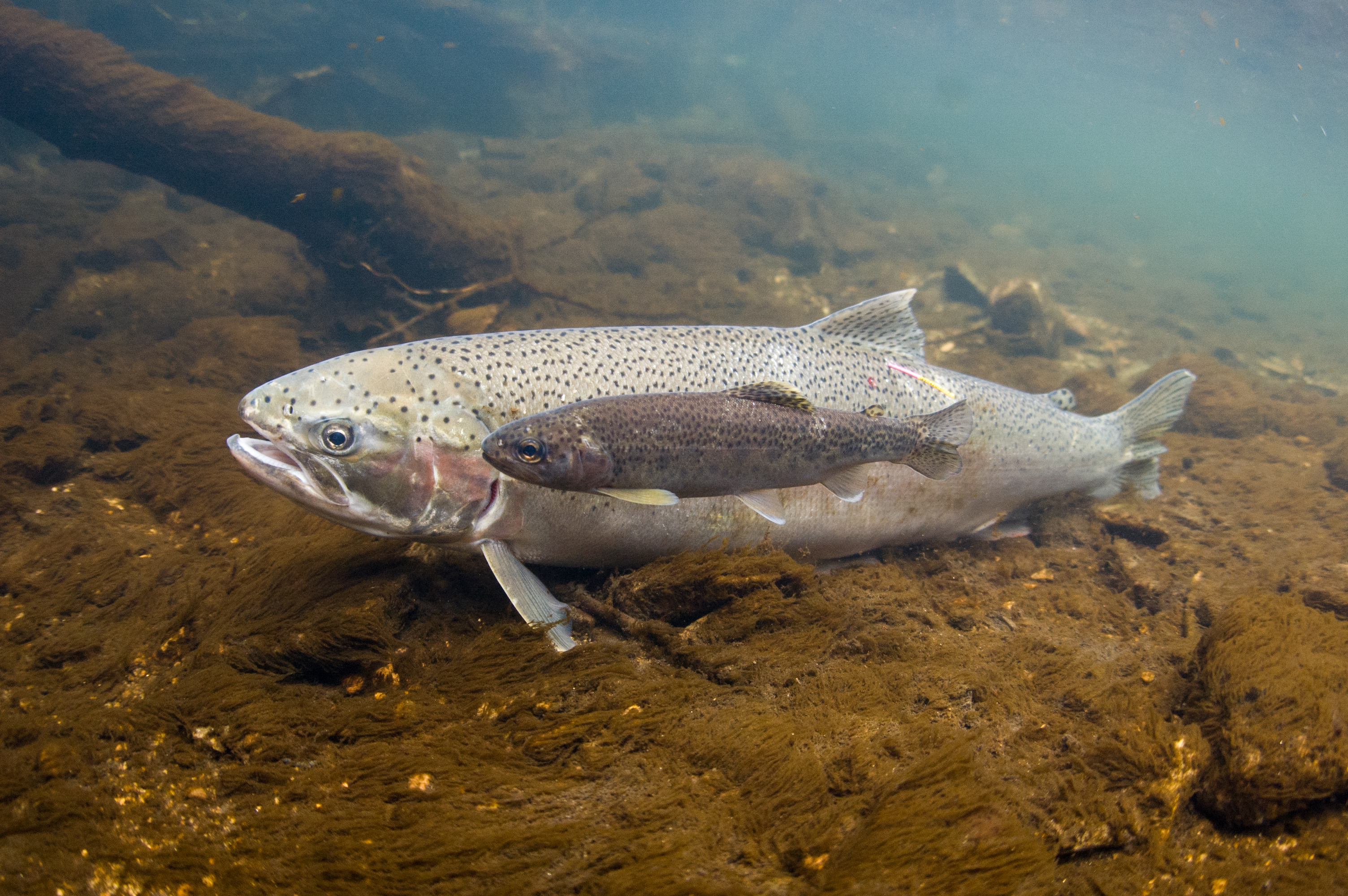
